# Mechanistic model of MAPK signaling reveals how allostery and rewiring contribute to drug resistance

**DOI:** 10.1101/2022.02.17.480899

**Authors:** Fabian Fröhlich, Luca Gerosa, Jeremy Muhlich, Peter K. Sorger

**Affiliations:** Laboratory of Systems Pharmacology, Department of Systems Biology, Harvard Medical School, 200 Longwood Avenue, Boston, MA 02115, USA; Genentech, Inc., South San Francisco, CA 94080

**Keywords:** allosteric interactions, rewiring, kinetic modeling, drug resistance, MAPK pathway

## Abstract

BRAF^V600E^ is prototypical of oncogenic mutations that can be targeted therapeutically and treatment of BRAF-mutant melanomas with RAF and MEK inhibitors results in rapid tumor regression. However, drug-induced rewiring causes BRAF^V600E^ melanoma cells to rapidly acquire a drug-adapted state. In patients this is thought to promote acquisition or selection for resistance mutations and disease recurrence. In this paper we use an energy-based implementation of ordinary differential equations in combination with proteomic, transcriptomic and imaging data from melanoma cells, to model the precise mechanisms responsible for adaptive rewiring. We demonstrate the presence of two parallel MAPK (RAF-MEK-ERK kinase) reaction channels in BRAF^V600E^ melanoma cells that are differentially sensitive to RAF and MEK inhibitors. This arises from differences in protein oligomerization and allosteric regulation induced by oncogenic mutations and drug binding. As a result, the RAS-regulated MAPK channel can be active under conditions in which the BRAF^V600E^-driven channel is fully inhibited. Causal tracing demonstrates that this provides a sufficient quantitative explanation for initial and acquired responses to multiple different RAF and MEK inhibitors individually and in combination.

**Highlights:**

- A thermodynamic framework enables structure-based description of allosteric interactions in the EGFR and MAPK pathways
- Causal decomposition of efficacy of targeted drugs elucidates rewiring of MAPK channels
- Model-based extrapolation from type I½ RAF inhibitors to type II RAF inhibitors
- A unified mechanistic explanation for adaptive and genetic resistance across BRAF-cancers

## INTRODUCTION

Eukaryotic signal transduction allows cells to regulate their growth, differentiation, and morphogenesis in response to external stimuli (Hunter, 2000; Ullrich & Schlessinger, 1990). In its reliance on receptor tyrosine kinase (RTK) autophosphorylation, assembly of signaling complexes on receptor tails, and activation of mitogen activated protein kinases (MAPKs; **Box 1**) signal transduction initiated by the binding of epidermal growth factor (EGF) to the EGF receptor (EGFR) is prototypical of growth- promoting signal transduction systems. The MAPK cascade comprises the RAF, MEK and ERK kinases, which regulate downstream factors such as ELK, ETS1 and AP1 transcription factors, as well as changes in cell motility and morphology (Lavoie *et al*, 2020). EGFR signaling has also been studied extensively using dynamical systems analysis (Starbuck & Lauffenburger, 1992; Kholodenko *et al*, 1999; Resat *et al*, 2003; Blinov *et al*, 2006; Chen *et al*, 2009; Gerosa *et al*, 2020) leading to better understanding of signal transduction in general as well as development of new modeling methods.

Oncogenic mutations are common in signal transduction networks and the V600E mutation in BRAF is an exemplar of these (Sanchez-Vega *et al*, 2018). In melanoma (Davies *et al*, 2002), thyroid cancer (Kebebew *et al*, 2007), colorectal cancer (Clarke & Kopetz, 2015), and other tissues, BRAF^V600E^ mutations cause constitutive activation of the MAPK pathway and oncogenic transformation. In cutaneous melanoma, inhibitors of the BRAF (BRAFi) and MEK (MEKi) kinases (e.g., vemurafenib and cobimetinib) are prototypical of highly effective targeted anti-cancer drugs (English & Cobb, 2002; Samatar & Poulikakos, 2014). A combination of BRAFi and MEKi is the current first-line treatment for metastatic melanoma (Sullivan & Flaherty, 2012) and frequently results in rapid tumor shrinkage. However, BRAF^V600E^ tumors usually develop resistance to RAFi/MEKi therapy within months to years, reducing long-term survival. The frequent and rapid rise of drug resistance in melanoma and the innate refractoriness of other MAPK-driven cancers to existing drugs has spurred extensive work aimed at understanding resistance mechanisms. Blocking the emergence of drug-resistant states is widely thought to be the key to achieving better patient outcomes with RAFi/MEKi drugs and precision oncology in general.

Resistance to MAPK inhibition occurs over a range of time scales. Adaptive resistance, which is reversible and does not involve acquisition or selection for mutations, can be observed within a few days of drug exposure (Fallahi-Sichani *et al*, 2017; Marin-Bejar *et al*, 2021; Oren *et al*, 2021). In cultured cells, adaptive resistance can last for months, giving rise to persister cells in which oncogenic BRAF signaling remains strongly inhibited but cells continue to grow, albeit more slowly than in the absence of drugs (Lito *et al*, 2012). In patients and in in cultured cells, acquisition of recurrent mutations, commonly in RTKs or components (or regulators) of the MAPK cascade, leads to reactivation of MAPK signaling and unrestrained cell growth (Shi *et al*, 2014; Long *et al*, 2014). The relationship between adaptive and acquired resistance is not fully understood and is an area of active investigation (Shaffer *et al*, 2017; Schuh *et al*, 2020). It is thought that DNA replication may be less faithful, or DNA damage responses less effective, in adapted than drug-naïve cells, leading to accumulation of resistance mutations (Russo *et al*, 2019; Shaffer *et al*, 2017; Schuh *et al*, 2020).

A paradox of the drug adapted state in BRAF^V600E^ mutant melanoma is that MAPK activity is known to be essential for proliferation of this cell type and yet oncogenic BRAF signaling remains strongly inhibited. Analysis of cell-average MAPK levels led to the suggestion that partial MAPK rebound (to ∼5% to 20% of the kinase activity in drug-naïve cells) is sufficient for cell survival and proliferation (Lito *et al*, 2012). However, more recent single-cell studies show that adapted cells experience sporadic MAPK pulses of ∼90 min duration and that these pulses are sufficient for cyclin D transcription and passage of a subset of cells into S phase (Gerosa *et al*, 2020). Pulses appear to arise from growth factors that act in an autocrine/paracrine manner by binding to EGFR and other RTKs expressed on persister cells. This finding raises a further question: how precisely can oncogenic MAPK signaling be repressed while receptor-mediated MAPK signaling remains active? The accepted explanation is that the cell signaling has become “rewired” in adapted cells (Ding *et al*, 2018; Lee *et al*, 2012; Wei *et al*, 2020).

In the absence of a new mutation, rewired networks are postulated to transmit or propagate oncogenic signals by different combinations or activity states of cell signaling proteins than drug-naïve networks. In some cases, rewiring is thought to involve a switch from one mitogenic pathway to another, from MAPK to PI3K-AKT signaling for example, but in drug resistant melanoma, the same MAPK components appear to be essential in the original and rewired states. More generally, rewiring is one of several concepts in translational cancer biology that are intuitively plausible but have not yet been subjected to quantitative, mechanistic modeling and analysis.

One way to gain deeper insight into rewiring at a mechanistic level is to perform the type of dynamical systems analysis that has previously proven effective in the study of RTK-MAPK signaling (Kholodenko *et al*, 1999; Rukhlenko *et al*, 2018; Kholodenko, 2015; Chen *et al*, 2009; Schöberl *et al*, 2009). This commonly involves constructing networks of ordinary differential equation (ODEs) to represent the precise temporal evolution of signal transduction networks under different conditions. ODEs are a principled way to represent cellular biochemistry in a continuum approximation and, with the addition of “compartments”, can also model the assembly of multi-protein complexes and transport between cellular compartments (Aldridge *et al*, 2006). In the case of the A375 melanoma cells used in this study, quantitative proteomics shows that proteins in the MAPK pathway are present at 10^2^ to 10^4^ molecules per cell (Gerosa *et al*, 2020), so continuum mass-action models represent an appropriate approximation (conversely, intrinsic noise is expected to be low).

Combinatorial complexity represents a substantial challenge to modeling even relatively restricted sets of signaling proteins. The presence of multiple reversible, post-translational modifications, protein-protein, and protein-small molecule interactions often makes the number of distinct biochemical species 10-1000 fold greater than the number of gene products (Faeder *et al*, 2005) (**Box 2**). Rule-based modeling was developed specifically to address this challenge and uses abstract representations of binding patterns and reactions to describe combinatorically complex networks in a compact programmatic formalism. Rules automatically generate ODE networks describing diverse types of reactions and molecular assemblies (Faeder *et al*, 2005; Hlavacek *et al*, 2006; Lopez *et al*, 2013) for subsequent model calibration and exploration.

An additional challenge in modeling MAPK signaling is that it involves allosteric regulation, in which the affinities of RAS, RAF and small molecules for each other are determined by protein conformation and oligomerization state. In conventional ODE modeling, a large number of parameters are necessary to describe the dependency of such affinities on states of assembly. However, protein-protein and protein- small molecule binding and unbinding does not consume energy and thermodynamic formalisms that impose energy conservation provide powerful means to constrain the number of binding parameters to a minimal, principled set (**Box 3**)(Ollivier *et al*, 2010; Sekar *et al*, 2016). The use of thermodynamics to derive kinetic rates was pioneered by Arrhenius (Arrhenius, 1889) and subsequently derived independently by Eyring, (Eyring, 1935), Evans and Polanyi (Evans & Polanyi, 1935), but it is only recently that practical approaches have emerged for using thermodynamic formalisms in reaction models (Gawthrop & Crampin, 2017; Honorato-Zimmer *et al*, 2015; Kholodenko, 2015; Klosin *et al*, 2020; Mason & Covert, 2018; Olivier *et al*, 2005; Rukhlenko *et al*, 2018; Gollub *et al*, 2021). Applications of these methods to signal transduction remain limited, in part because of the complexity of relevant models, but Kholodenko and colleagues have pioneered the application of thermodynamic balance to MAPK signaling (Rukhlenko *et al*, 2018).

Model calibration and non-identifiability represents a final challenge in modeling networks of readily reversible reactions. Model calibration (estimating parameter values that minimize the deviation from experimental data) is compute-intensive (Fröhlich *et al*, 2017) and even after calibration, parameters can assume wide ranges, a property known as non-identifiablity (Kreutz *et al*, 2012; Raue *et al*, 2011; Kreutz *et al*, 2012; Chis *et al*, 2011; Wieland *et al*, 2021). When models are combinatorically complex and non-identifiable it can be difficult to quantify fluxes, explain how signaling state arise and trace how species of interest are created by upstream reactions and consumed downstream. This complicates the quantification of signal propagation through the reaction network, a prerequisite for the investigation of concepts of such as network rewiring.

In this paper we described a second-generation MAPK Adaptive Resistance Model (**MARM2.0**) that seeks to explain the rewiring of EGFR/MAPK signaling occurring in drug adapted BRAF^V600E^ melanoma cells. MARM2.0 builds on a large body of structural, biochemical and theoretical work on EFGR/MAPK signaling and feedback regulation (Haling *et al*, 2014; Hatzivassiliou *et al*, 2013; Lito *et al*, 2012, 2013; Poulikakos *et al*, 2010; Solit *et al*, 2006; Yao *et al*, 2015) and is constructed using rule-based modeling in PySB with thermodynamic balance. By developing a new approach to causal tracing, we show how rewiring alters the organization and amplification/attenuation characteristics of multiple reaction channels operating in parallel in the MAPK cascade. We find that, in addition to the well-known differential sensitivity of oncogenic RAF monomers and wild-type dimers to RAFi, there exists a similar, less characterized differential sensitivity of MEK to MEKi based on whether the signal arises from BRAF^V600E^ or wild-type RAF. Together with a time-scale separation between signal transduction and transcriptional feedback, this generates a drug adapted state in which BRAF^V600E^ is inhibited but a MAPK cascade involving many of the same components can be activated by RTK ligands or mutation of proteins such as NRAS.

## RESULTS - TEXT BOXES 1 TO 3

### Box 1. The MAPK signaling pathway

The core of the MAPK pathway is a three-enzyme cascade comprising RAF-MEK-ERK kinases (HUGO: ARAF/BRAF/RAF1, MAP2K1/MAP2K2, and MAPK1/MAPK3) that transduces signals from extracellular stimuli, most commonly growth factors and receptor tyrosine kinases (RTKs) (Lavoie *et al*, 2020). Three-enzyme cascades involving closely related kinases also transmit signals from cytokines and their receptors. Driving oncogenic mutations are found in multiple components in or upstream of the MAPK pathway (Burotto *et al*, 2014), commonly KRAS (G12C/D/V, G13C/D), NRAS (Q61H/K)(Prior *et al*, 2012), BRAF (V600E/K) and less commonly MEK and ERK (Gao *et al*, 2018). BRAF^V600E^ or closely related mutations (e.g., BRAF^V600K^) are found in ∼50% of cutaneous melanomas and RAF/MEK therapy is the first line treatment option for BRAF-mutant metastatic melanoma (Flaherty *et al*, 2012). BRAF mutations are also found in ∼10% of colorectal cancers and several other tumor types (Davies *et al*, 2002), but RAF/MEK therapy is rarely effective in these settings.

Binding of growth factors to RTKs induces their intracellular auto-phosphorylation, followed by association of SH2 and SH3-containing proteins with phosphorylated tyrosine residues on receptor tails. Subsequent signalosome assembly involves adaptor proteins such as GRB2, enzymes that modify second messengers such as PI3Ks, and guanine nucleotide exchange factors (GEFs) such as SOS1 (Lemmon & Schlessinger, 2010). GEFs convert one or more of the N, K, and H RAS GTPases (depending on cell type) into the active GTP-bound form, and GTP-bound RAS then activates the ARAF/BRAF/RAF1 kinases by recruiting them to the plasma membrane and inducing their dimerization. BRAF/RAF1 homo- and heterodimers are the primary mediators of MEK phosphorylation (ARAF has low kinase activity). Phosphorylated and active MEK then phosphorylates ERK on two proximate residues. Both phosphorylation steps are potentiated by the assembly of multi-protein complexes involving 14-3-3 and KSR scaffolding proteins (Lavoie & Therrien, 2015). Active ERK phosphorylates transcription factors, cytoskeletal proteins, and other kinases and is the proximate functional output of the MAPK cascade. Changes in the levels or activities of proteins such as DUSP4/6 phosphatases, which remove activating phosphorylation modifications, and SPRY2/4 proteins, which sequester GRB2, as well as inhibitory phosphorylation of EGFR, SOS1 and CRAF act as negative-feedback mechanisms and enforce homeostatic control over MAPK activity.

### Box 2. Drugs targeting MAPK kinases

Multiple small molecule inhibitors targeting individual MAPK kinases are FDA approved but combinations of RAF and MEK inhibitors are the most widely used clinically. A subtle relationship exists between the mechanism of action of these drugs, kinase conformation, and formation of mutli-protein complexes. In the absence of upstream stimuli, RAF kinases are present in cells as monomers but activation by RAS-GTP causes dimerization. Some activating BRAF mutations (Yao *et al*, 2015) and splice variants (Poulikakos *et al*, 2011) also promote dimerization, but BRAF^V600E/K^ kinases are constitutively activated without requiring dimerization. Whether RAF is present in monomer, heterodimer or homodimer forms profoundly influences the enzyme’s sensitivity to inhibition (Yao *et al*, 2015). The FDA approved RAF inhibitors vemurafenib, dabrafenib, and encorafenib are ATP-competitive type I**½** kinase inhibitors (Roskoski, 2016) that preferentially bind to the alpha-C helix-out, DFG-in conformation assumed by BRAF^V600E/K^; this state differs from the alpha-C helix-in (and DFG-in) state found in activated wild-type RAF (Karoulia *et al*, 2017). Whereas binding of type I**½** BRAF inhibitors to BRAF^V600E/K^ is inhibitory, binding to wild type RAF monomers promotes kinase dimerization and activation, leading to amplification of MAPK signaling, a phenomenon termed paradoxical activation (Hall-Jackson *et al*, 1999; Poulikakos *et al*, 2010; Hatzivassiliou *et al*, 2010). To prevent this, “paradox breaker” RAF inhibitors such as PLX8394 have been developed (Tutuka *et al*, 2017; Yao *et al*, 2019; Zhang *et al*, 2015). These are type I**½** inhibitors that, by virtue of locking the R506 side-chain in the out conformation, do not promote dimerization (Karoulia *et al*, 2017). Both regular and paradox breaker type I**½** inhibitors have a lower affinity for the 2nd protomer in a RAF dimer, which typically assumes the inactive alpha-C helix- in, DFG-out conformation. Thus, the structural differences between monomers and dimers (rather than differences in the ATP binding pocket) are the basis of the selectivity of clinically approved RAF inhibitors for cells transformed by BRAF mutant kinases. However, the inability of type I**½** inhibitors to fully inhibit homo- and hetero-dimer RAF kinases is also a primary mechanism of drug resistance in cancers with sustained RAS-GTP signaling; one well established example is EGFR-driven signaling in BRAF^V600E/K^ colorectal cancer. In contrast, so-called “panRAF” Type II inhibitors, such as the Phase 1 compound LY3009120 (Peng *et al*, 2015) and preclinical compound AZ-628 (Noeparast *et al*, 2018), bind RAF in the alpha-C helix-in, DFG-out conformation and can, thus, bind both RAF protomers with similar potency. These inhibitors can achieve more complete MAPK suppression but appear to cause additional toxicity, presumably by interfering with MAPK activity in non-cancer cells. Multiple type II inhibitors are currently under clinical investigation for solid tumors (Yen *et al*, 2021), including melanoma, but, so far, none have been approved for use in humans.

FDA approved MEK inhibitors such as cobimetinib, trametinib and binimetinib, are type III non-ATP competitive (allosteric) inhibitors that lock the MEK kinase in a catalytically inactive state, limit movement of the activation loop, and decrease phosphorylation by RAF (Wu & Park, 2015). These MEK inhibitors are more potent at preventing ERK activation by BRAF^V600E/K^ than by RAF acting downstream of mutant RAS (Lito *et al*, 2014; Hatzivassiliou *et al*, 2013) or RTKs (Gerosa *et al*, 2020). The reasons for this are not fully understood but are thought to include the lower affinity of MEK inhibitors for phosphorylated as compared to unphosphorylated MEK and differences in RAF-MEK binding (Hatzivassiliou *et al*, 2013; Pino *et al*, 2021).

### Box 3. Thermodynamic description of conformational states in rule-based modelling

Changes in protein assembly and conformation, often mediated by post-translational modification, are the structural basis for much of signal transduction. For example, generating the active conformation of CRAF requires both N-terminal phosphorylation and association with a second RAF family member to stabilize the active state. Because formation of protein-protein interactions does not consume energy, a strict relationship exists between conformation and binding affinity (Tsai & Nussinov, 2014): when binding increases the stability of a specific conformational state, that state will also have higher binding affinity for its interacting partner. Since this relationship is transitive, binding affinities can be coupled through conformational states, giving rise to long-range, higher-order dependencies in oligomeric complexes. Such higher-order dependencies can create ultrasensitive responses, which are often involved in cell fate decisions or homeostasis.

A conformational state is defined by a specific local minimum in the Gibbs free energy landscape. The relative stability of a conformational state *S* can be expressed as free energy difference *ΔG*_*c*_ with respect to a reference state *S*_0_. Stabilizing or destabilizing conformational states is equivalent to changes in this free energy difference (i.e., *ΔΔG*_*c*_). Similarly, binding reactions can be characterized by the difference *ΔG*_b_ between the Gibbs free energies of binding educts and binding products, which is proportional to the logarithm of their dissociation constant *K*: *ΔG*_b_ = −*RTlog*(*K*), where R is the gas constant and T is the temperature. Energy conservation guarantees that a ligand (L)-induced changes to the free energy of a conformational state *S* (*ΔΔG*_*c*_) is equal to the difference *ΔΔG*_b_ in the affinity of L for *S* as compared to *S*_0_. This equilibrium description can be extended to dynamic behavior by means of the Arrhenius Equation (Arrhenius, 1889), which defines reaction propensities according to the free energy of the transition state (Sekar *et al*, 2016). Such an energy-based formulation enforces Wegscheider-Lewis cycle conditions on kinetic parameters (Wegscheider, 1911), ensuring detailed balance for equilibrium states, but also constraining dynamics of non-equilibrium processes. By ensuring energy conservation, the effective number of parameters needed to describe multimeric oligomerization processes is reduced (Kholodenko, 2015) and powerful constraints are placed on the structures of models describing species that adopt multiple conformational states.

Energy conservation provides a natural framework for the specification of structure-based kinetic models that include allosteric interactions (Rukhlenko *et al*, 2018) and has been incorporated into a rule-based modeling form as energy-BioNetGen (eBNG)(Sekar *et al*, 2016). In eBNG, allosteric interactions are encoded using energy patterns that permit specification of *ΔΔG*_b_. For example, a kinetic model for the binding of RAF inhibitors (RAFi in text, I in figure) to RAF kinases (RAF in text, R in figure) (**Figure Box 1A**) can be constructed using one rule for RAF dimerization (turquoise) and another for drug binding to RAF (black), which generates 12 reversible reactions (**Figure Box 1B**). Allostery for drug binding to the 1^st^ or 2^nd^ protomer of a RAF dimer is imposed using the thermodynamic factors *f* (orange) and *g* (purple), which change *ΔΔG*_b_ via two energy patterns. The contribution of these thermodynamic factors to kinetic rates is exemplified by the relationship between Gibbs free energies and rate constants for RAF dimerization that are RAFi-dependent (**Figure Box 1C**; no RAFi, black; one RAFi, orange; two RAFi purple). The parameter ϕ, controls whether *ΔΔG*_b_ influences educt states (ϕ = 0) or product states (ϕ = 1, depicted in C) or a mixture (0< ϕ <1). Using PySB, all 12 reactions depicted in **Figure Box 1B** can be specified using two rules and four energy-patterns (**Figure Box 1D)**. Thus, PySB code automatically generates symbolic reaction rates that parameterize the reaction network according to allosteric effects whose magnitudes are set by the thermodynamic factors *f* and *g* (**Figure Box 1E).** In this way, models of complex drug-protein interactions, such as resistance mediated by formation of RAF dimers, can be easily parameterized in terms of the baseline equilibrium constant for RAF dimerization (*K_RR_*). We illustrated this by simulations with *f*=0.001 and *g*=1000 (**Figure Box 1F**) which represent a type I**½** RAF inhibitor that avidly binds the 1^st^ RAF protomer but has a 10^6^-fold lower affinity for the 2^nd^ protomer in a RAF dimer.

**Figure Box1:**
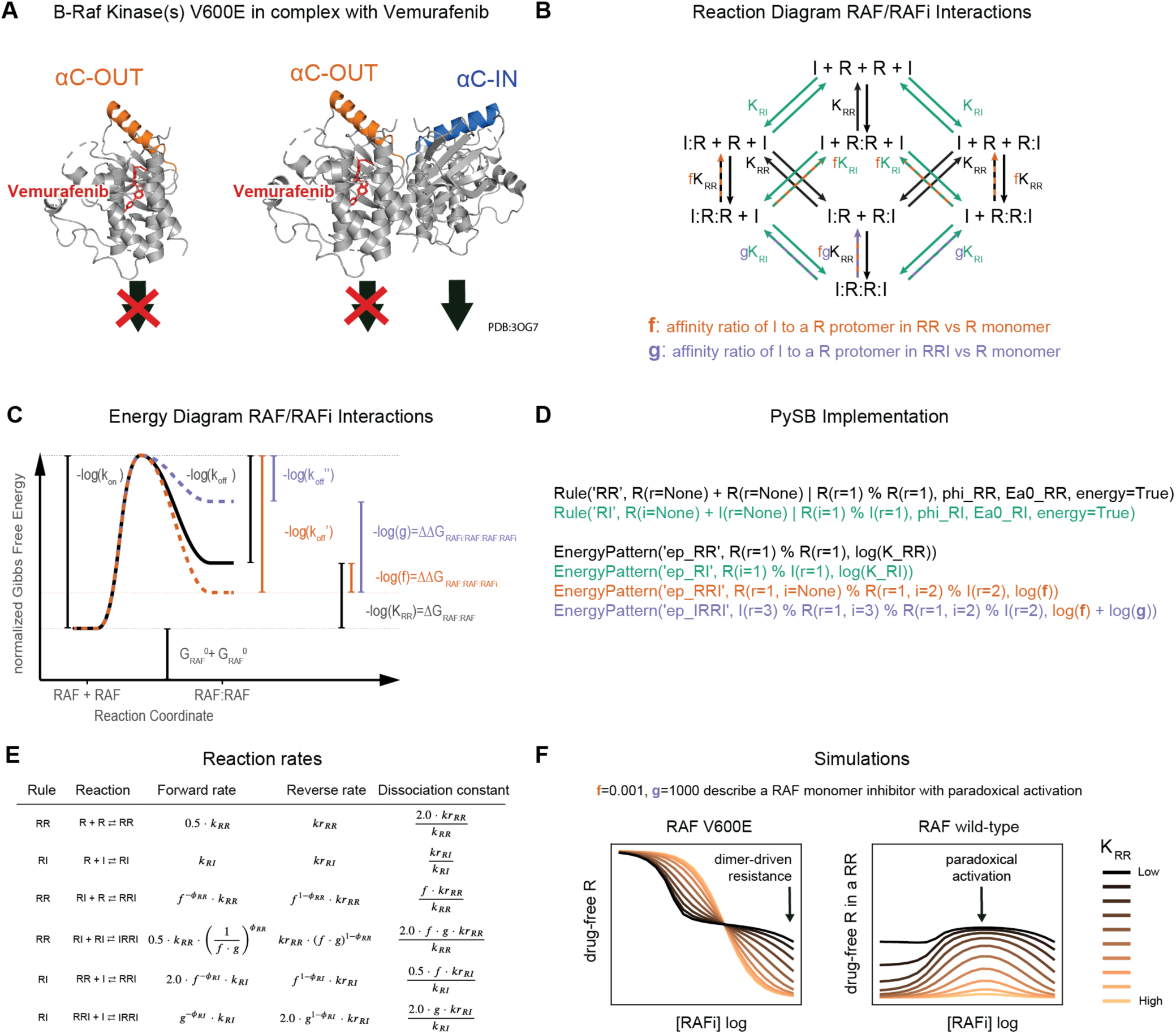
Thermodynamic Model of RAF-RAFi interactions. **(A)** Protein structures of monomeric and dimeric BRAF^V600E^ protomers bound to vemurafenib **(B)** Binding diagram for RAF and RAFi molecules. Formulas next to reaction arrows indicate the dissociation constants of the respective reactions. Arrow color indicates type of reaction (black: RAF dimerization, turquoise: RAFi binding). Dashed line color indicates the thermodynamic parameters that modulate the respective reactions (orange: f, purple: g). **(C)** Illustration of relationship between Gibbs free energies and kinetic rates for RAF dimerization. Modulation of kinetic rates through a context specific energy patterns that depends on the number of bound RAFi molecules is indicated in orange (one RAFi bound, parameter f) and purple (two RAFi bound, parameter g). Energies are normalized by the factor 1/RT, where R is Gas constant and T is the temperature. The diagram shows the specific situation of *ϕ* = 1 where only reaction product stability is modulated. (**D**) PySB code to define the rules and energy-patterns that describe the diagram in B. **(E)** Table of context dependent forward and reverse reaction rates. k is the binding rate, kr is the unbinding rate, with corresponding pysb rule indicated as subscript. **(F)** Model simulations for different values of K_RR_ with f=0.001 and g=1000

## MAIN RESULTS

### A Structure-Based Model of EGFR and ERK Signaling

The MAPK signaling cascade (**Box 1**) and its immediate regulators constitute no more than two dozen unique gene products, but the binding of these proteins to each other gives rise to a remarkably large number of molecular species, many of which have distinct activities. Moreover, the complexity of the MAPK cascade increases substantially when we consider states that are bound and unbound to drugs. For example, BRAF/CRAF can exist in monomeric, homo- and heterodimeric forms, with either one or two subunits bound to RAFi, each with or without RAS-GTP bound as an activator. Drug binding occurs preferentially to some BRAF oligomers and not others (**Box 2**), and can strongly influence association with upstream and downstream factors. To recapitulate the responses of cells to RAFi in a mechanistic computational model, it is necessary for the allosteric interactions that control association of RAS, RAF and RAFi to be described in detail (Rukhlenko *et al*, 2018).

To accomplish this, we generated a compartmentalized ODE model of MAPK signaling (the MAPK Adaptive Resistance Model MARM2.0) that extends a related model (MARM1.0) used in an experimental study we recently published (Gerosa *et al*, 2020) that uses modeling as an explanatory tool but does not involve any model analysis. Such analysis is the focus of the current paper and its updated model. MARM2.0 was calibrated using data described in Gerosa *et al*. with the addition of drug-response data that is unique to the current study. Moreover, both MARM1.0 and MARM2.0 build on an earlier model of RAF-RAFi interaction developed by Kholodenko (Kholodenko, 2015), but with the inclusion of more proteins and complexes. Model expansion was greatly facilitated by the use of rule-based BNG models in the domain-specific Python language PySB (Blinov *et al*, 2004; Lopez *et al*, 2013). More specifically, MARM1.0 & 2.0 extend the RAF-MEK-ERK model of Kholodenko with the addition of upstream activation and multiple feedback mechanisms relevant to acquired resistance to RAF inhibitors (Lito *et al*, 2012) and a more detailed description of MAPK enzymes (**Figure 1A)**. Compared to MARM1.0, MARM2.0 is compartmentalized (compartments: *extracellular space*, *plasma membrane*, *cytoplasm* and *endosomal membrane*), it adds EGFR-CBL interaction and endosomal recycling, and includes mRNA species in the description of transcriptional feedback control. In total, MARM2.0 involves 17 distinct molecular species: eleven proteins, three mRNA species and three small molecule inhibitor classes. Proteins include EGFR, BRAF, CRAF, MEK and ERK, the dual specificity phosphatase DUSP, guanine nucleotide exchange factor SOS1, GTPase RAS, E3 ubiquitin ligase CBL, adaptor protein GRB2, and RTK negative regulator SPRY (ellipses in **Figure 1A**). RAFi, panRAFi and MEKi, (depicted as colored circles and rounded boxes in **Figure 1A**) are optionally present and values for kinetic and energetic parameters can be set so that the inhibitors can correspond to any of ten different small molecules that are used as human therapeutics or pre-clinical tools. These comprise the RAFi compounds vemurafenib, dabrafenib, PLX8394, the panRAFi compounds LY3009120 and AZ628, and MEKi compounds cobimetinib, trametinib, selumetinib, binimetinib and PD0325901.

**Figure 1:**
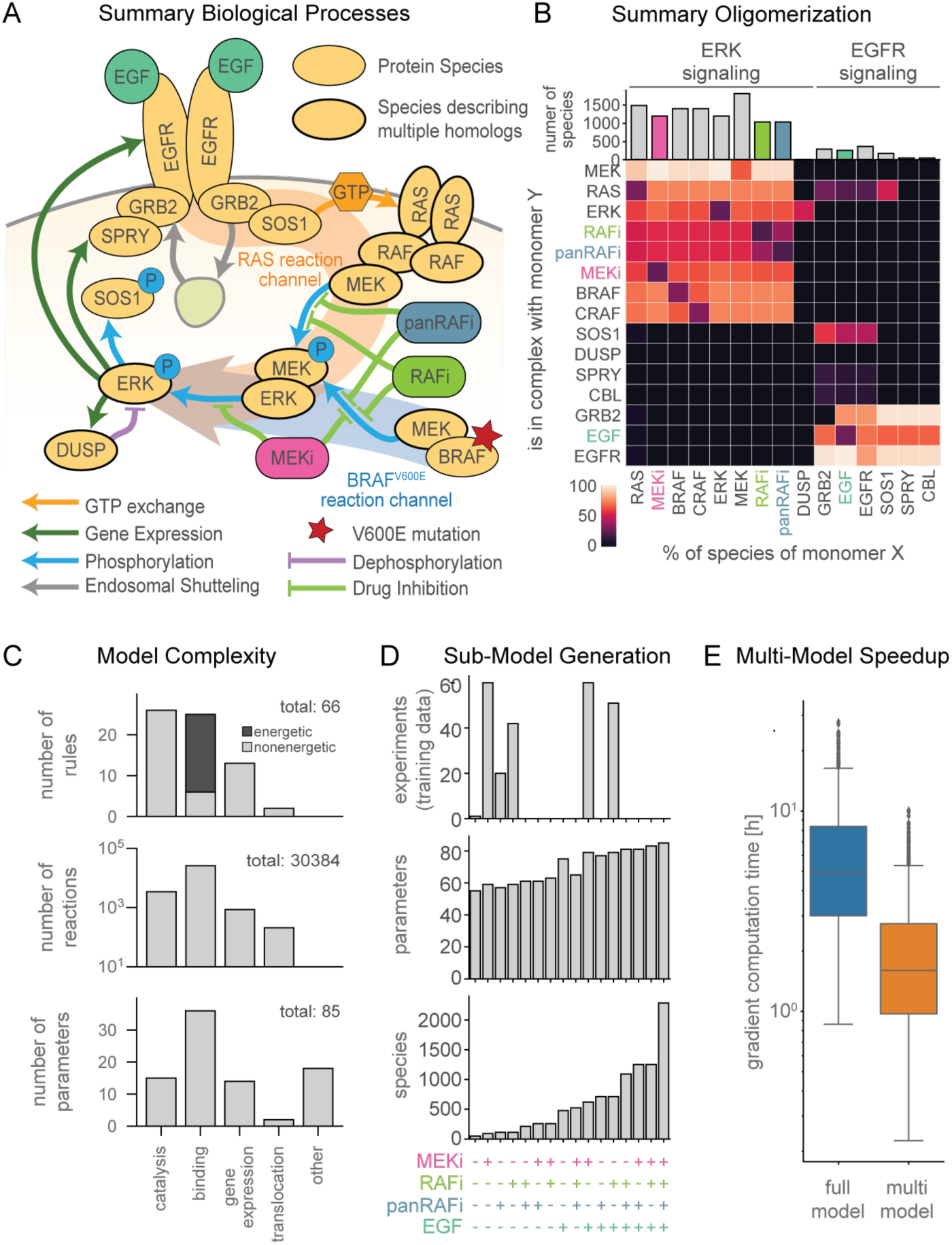
Thermodynamic model of EGFR and ERK signaling. **(A)** Schematic overview of processes described in the model. **(B)** Summary of model species and oligomerization in the model. Coloring of tiles indicates percentage with respect to total of monomer species (per row). Columns for the drug and growth factor perturbations RAFi, panRAFi, MEKi and EGF are highlighted according to the respective color in A. **(C)** Statistics of model rules, reactions and parameters. Catalysis includes (de-) phosphorylation, GTP- exchange and (de-)ubiquitination. Other parameters include initial conditions and scaling factors and background intensities. **(D)** Number of experiments and sizes of respectively resized models according to the multi-model optimization scheme. A plus on the bottom indicates that the respective perturbation was applied in the corresponding experiment, color as in A/B. **(E)** Comparison of gradient computation time for the full-model and multi-model optimization approach.

To maintain model tractability we lumped together paralogs, combined phosphorylation sites having similar functions, and simplified other aspects of EFGR regulation, which exhibits particularly high combinatorial complexity (Blinov *et al*, 2006). MARM2.0 nonetheless has over 15,000 biochemical reactions, illustrating how transient binding among a few kinases, their regulators, and inhibitory drugs generates an elaborate biochemical network. With respect to paralogs, we made the following assumptions: “RAS” stands in for KRAS, NRAS, and HRAS, “MEK” for MAP2K1 and MAP2K2, “ERK” for MAPK1 and MAPK3, “DUSP” for DUSP4 and DUSP6, and “SPRY” for SPRY2 and SPRY4 (lumping of paralogs is depicted in **Figure 1A** by thick outlines). This is equivalent to assuming that all paralogs have the same kinetic rate constants. In some cases, paralogs are known to be very similar (e.g., MAPK1, MAPK3) but in other cases they are functionally distinct (e.g, KRAS, NRAS and HRAS). The three RAS paralogs are expressed at similar levels in A375 and we did not distinguish among them because we do not yet have relevant training data. However, MARM could easily be modified for future studies that focus on differences between RAS species. We did not lump BRAF and CRAF into a single RAF species due to the unique role that BRAF^V600E^ plays as an oncogene; ARAF was omitted due to its low kinase activity. We also lumped together multi-site phosphorylation of EGFR (on Y1068, Y1086, Y1173, etc.), MEK (MAP2K1: S218, S222; MAP2K: S222, S226) and ERK (MAPK1: T185, Y187; MAPK3: T202, Y204) as single post translational modifications for each protein. The underlying phosphorylation reactions were implemented as two-step reactions comprising substrate binding and phosphorylation steps. Finally, mRNA species were included for DUSP, EGFR and SPRY to model transcriptional feedback with distinct, lumped translation rates for each species (depicted by dark green arrows in **Figure 1A**). This made it possible to calibrate models on time-course and dose-response transcriptomic data.

To model RTK-induced MAPK activation we focused on EGFR autophosphorylation at Y1068, Y1086 and Y1173, which creates GRB2 binding sites (Batzer *et al*, 1994) as well as EGFR ubiquitination by CBL (Alwan *et al*, 2003) and subsequent endocytosis and recycling. EGFR endocytosis and recycling rates were dependent on EGFR levels, as previously described (Starbuck & Lauffenburger, 1992; Resat *et al*, 2003). The “addition” of EGF to MARM2.0 promotes EGFR dimerization and trans- phosphorylation, recruitment of GRB2:SOS1 complexes to phospho-tyrosine residues on receptor tails and consequent GTP loading and activation of RAS. Receptors are then subjected to endocytosis leading to either their degradation or recycling. GTP-loaded RAS (RAS-GTP) promotes RAF dimerization and initiates the RAF-MEK-ERK (MAPK) cascade (**Box 2**). When BRAF^V600E^ is present, it constitutively phosphorylates MEK in the absence of upstream signals. Phosphorylated MEK (pMEK) phosphorylates ERK (pERK), which indirectly upregulates expression of proteins that act as negative regulators of RTK signal transduction (these intermediate steps are represented as lumped reactions). Multiple negative regulatory mechanisms are known and we modelled four of them. Three involved transcriptionally- mediated changes in protein abundance for (i) EGFR itself, (ii) DUSP, which antagonize ERK signaling by dephosphorylating the T and Y residues in the T-Y-X motif in the ERK activation loop (Saha *et al*, 2012; Corbalan-Garcia *et al*, 1996) and (iii) SPRY, which has multiple biochemical activities, among which we modeled sequestration and inactivation of GRB2 (Lao *et al*, 2006, 2). We also modeled the phosphorylation-dependent inhibition of SOS1 binding to GRB2 and acquisition of a 14-3-3 docking site, which sequesters the protein in an inactive conformation (Corbalan-Garcia *et al*, 1996; Kamioka *et al*, 2010). SOS1 is phosphorylated on S1134 and S1161 sites by RSK, which is transcriptionally and post-translationally activated by ERK, but we represented this with a single pERK dependent phosphorylation reaction.

MARM 2.0 includes 66 rules and 85 free parameters (kinetic rates, energies, scaling factors, etc.; total 109 free parameters when instantiating MARM2.0 for all of the 10 small molecules). Six rules described transcript turnover, 7 protein turnover, 22 phosphorylation, 25 binding and 3 sets of 2 rules each described GTP/GDP exchange, ubiquitination, and translocation between cellular compartments (**Figure 1C**). For example, the binding rule “*Rule(’BRAF_and_uMEK_bind_and_dissociate’, BRAF(mek=None) + MEK(phospho=’u’, raf=None) | BRAF(mek=1) % MEK(phospho=’u’, raf=1), …*)” describes binding of BRAF to unphosphorylated MEK (uMEK), a prerequisite for MEK phosphorylation. Binding requires MEK to be unphosphorylated (*phospho=’u’*), but does not specify any dependence on RAS, BRAF, CRAF or RAFi. Implementation of PySB rules generated >2,200 molecular species and >30,000 biochemical reactions with most proteins participating in >1000 species, a reflection of the combinatorial complexity described above. Binding rules accounted for >85% of all reactions in the model (25,922 of 30,384 reactions total) and > 75% (19/25) of these binding rules were formulated as “energetic rules” with binding affinities expressed in terms of normalized Gibbs free energy differences (ΔG; **Box 3**). Binding and unbinding rates were then computed according to the Arrhenius law. To facilitate programmatic model formulation within an energetic framework, we implemented support for the eBNG framework (Hogg, 2013; Harris *et al*, 2016) in PySB. This enabled specification of allosteric interactions using differences in free energy differences (ΔΔG, **Box3**), which is a principled way of establishing context dependent binding and unbinding rates (with the balance encoded by the parameter ϕ).

### ODE Description of ERK Pulsing Enabled Use of Population Average and Perturbational Experiments to Describe the Behavior of Single Cells

Imaging studies have established that the A375 BRAF^V600E^ melanoma cell line used in this study enters a seemingly steady-state drug-adapted condition within 24 hours of exposure to RAFi and/or MEKi (Gerosa *et al*, 2020). Data were therefore collected at this time point or subsequently, and model simulations included a pre-equilibration step. Once adapted to RAFi, BRAF^V600E^ melanoma cells experience transient pulses of ERK activity at irregular intervals, consistent with a stochastic regulatory mechanism (Gerosa *et al*, 2020). In principle, BNG/PySB models can be instantiated as stochastic, agent-based systems to represent such stochastic fluctuations (Sneddon *et al*, 2011). However, the reactions in MARM2.0 involve sufficiently abundant proteins (∼10^2^ to 10^6^ copies per cell) that intrinsic stochasticity is not expected to arise spontaneously. Thus, the irregular pulsing by drug adapted A375 cells appears to originate not in the noise of intracellular reactions, but instead in the spatially restricted release of growth factors acting in an autocrine and paracrine manner (Gerosa *et al*, 2020). In the absence of better understanding of these extracellular processes, they are difficult to represent computationally. Moreover, calibration of stochastic models is substantially more difficult than for deterministic models (Fröhlich *et al*, 2016).

Fortunately, experiments showed that addition of any of several different exogenous growth factors to RAFi- or MEKi-adapted cells generates synchronous ERK pulses having the same dynamics and drug sensitivities as asynchronous pulses arising spontaneously (Gerosa *et al*, 2020). Because single cells are much more similar to each other during ligand-induced than spontaneous pulsing, induced pulses are more amenable to characterization using standard transcriptional profiling and protein mass spectrometry methods. A further advantage is that synchronous pulses can be modeled at the population level by an ODE-model that is a reasonable simulacrum of single cell biology. In the current work, we used data from pulses generated by growth factors to provide insight into spontaneous pulses; as a consequence, we focused only on mechanisms downstream of receptor activation. Future work will be required to understand the origins and spatial distributions of ligands in the micro-environment of drug adapted cells undergoing asynchronous and spontaneous pulsing.

To further constrain MARM2.0, we used targeted proteomics with calibration peptides to measure the absolute abundances of all 11 protein species and two phospho-proteins; data were collected at five vemurafenib concentrations yielding 55 data points for model calibration. In addition, we extracted relative abundances for 3 mRNA species from genome-wide transcript profiling performed at 8 vemurafenib concentrations and 7 timepoints following EGF stimulation (yielding 45 calibration data points). Immunofluorescence imaging of pERK and pMEK provided the greatest amount of data (847 data points) and involved 234 different experimental conditions each involving a different concentration of one or more of the following perturbations: EGF, RAFi, panRAFi or MEKi. Imaging data had single cell resolution but population averages were used for model calibration, since we aimed to model the behavior of an average single cell. Training data was complimented with 2,209 immunofluorescence data points in 1,647 conditions for model validation, which are described in greater detail below.

### Rule-Based Modeling enables Efficient Calibration through Multi-Model Optimization

To calibrate MARM2.0 on experimental data, we used gradient-based numerical optimization, which performs well for large models (Villaverde *et al*, 2019). Optimization is nonetheless challenging for a model with as many reactions as MARM2.0: weighted least squares minimization of an objective function required simulation for each of the 234 training conditions for every evaluation of the objective function, and this took minutes to perform. Optimization required hundreds of evaluations of the objective function and its derivatives, resulting in calibration runtimes on the order of weeks to months even on a cluster computer. However, we found that, by exploiting patterns in the perturbational data it was possible to substantially reduce the number of species in a condition-specific manner, accelerating calibration (Fröhlich *et al*, 2019; Städter *et al*, 2021). In our calibration dataset, 122 conditions involved one perturbation (RAFi, panRAFi or MEKi individually), 111 conditions involved two perturbations (RAFi or MEKi followed by addition of EGF) and only one involved no perturbation, (**Figure 1D**, top). In the absence of a perturbing agent, all model species involving that agent (e.g., RAF bound to RAFi, **Figure 1B**) as well as a subset of downstream species (e.g., pEGFR activated by EGF) have zero concentrations and need not be modelled. To automatically generate, compile and track sub-models omitting zero concentration species for a diverse range of perturbations, we created routines that exploited the programmatic features of PySB (Lopez *et al*, 2013) and BNGL network generation (Blinov *et al*, 2004) (see MultiModelFitting in Material and Methods). This yielded models having an average of 1.5 times fewer parameters than MARM2.0 itself (55-83 parameters compared to 85) (**Figure 1D**, middle) and up to 45-fold fewer species (50-1253 species compared to 2284) (**Figure 1D**, bottom). Multi-model objective calibration was performed using pyPESTO (a python reimplementation of the Parameter EStimation Toolbox; (Stapor *et al*, 2018)) allowing consistent generation of a full model based on calibration of sub- models; this is an exact approach that does not reduce the accuracy of the objective function or gradient evaluation. Overall, we found that using PySB to match model structure to data structure reduced median gradient evaluation time ∼3-fold (from 5h to 1.60h on a single compute core; **Figure 1F**), which for MARM2.0 extrapolated to a reduction of ∼2 weeks in wall-time and ∼38 years in CPU time (using 10^3^ cores with 5 days wall-time). Since multiple rounds of model refinement and calibration were necessary over the course of the current work, a three-fold improvement in calibration time had a major impact. We expect that multi-model objective calibration will be broadly useful with other models involving perturbational datasets.

Following calibration, MARM2.0 quantitatively captured the effects of RAFi and MEKi treatment on baseline pERK levels in the drug adapted state and during transient EGF stimulation. Relatively few parameters converged on unique values (**Figure S1)** due to the known non-identifiability of biochemical models having explicit forward and back reactions, (Gutenkunst *et al*, 2007) as well as incomplete convergence of the optimizer due to limitations in the computational budget. We therefore used parameter sets from the 5% of optimization runs having the lowest value of the objective function (50 parameter sets) to generate a set of dynamical trajectories that estimated the impact of parametric uncertainty on simulations. For the great majority of data points (87.4%) we found that 80% of simulated trajectories fell within experimental error bounds (**Figure 2, S2**), demonstrating good agreement between the calibrated model with experimental data. This does not constitute a rigorous quantification of parameter uncertainty (Fröhlich *et al*, 2014), but does account for correlation in parameter values (Eydgahi *et al*, 2013) and was the only practically applicable approach given the number of parameters and species in MARM2.0.

**Figure 2:**
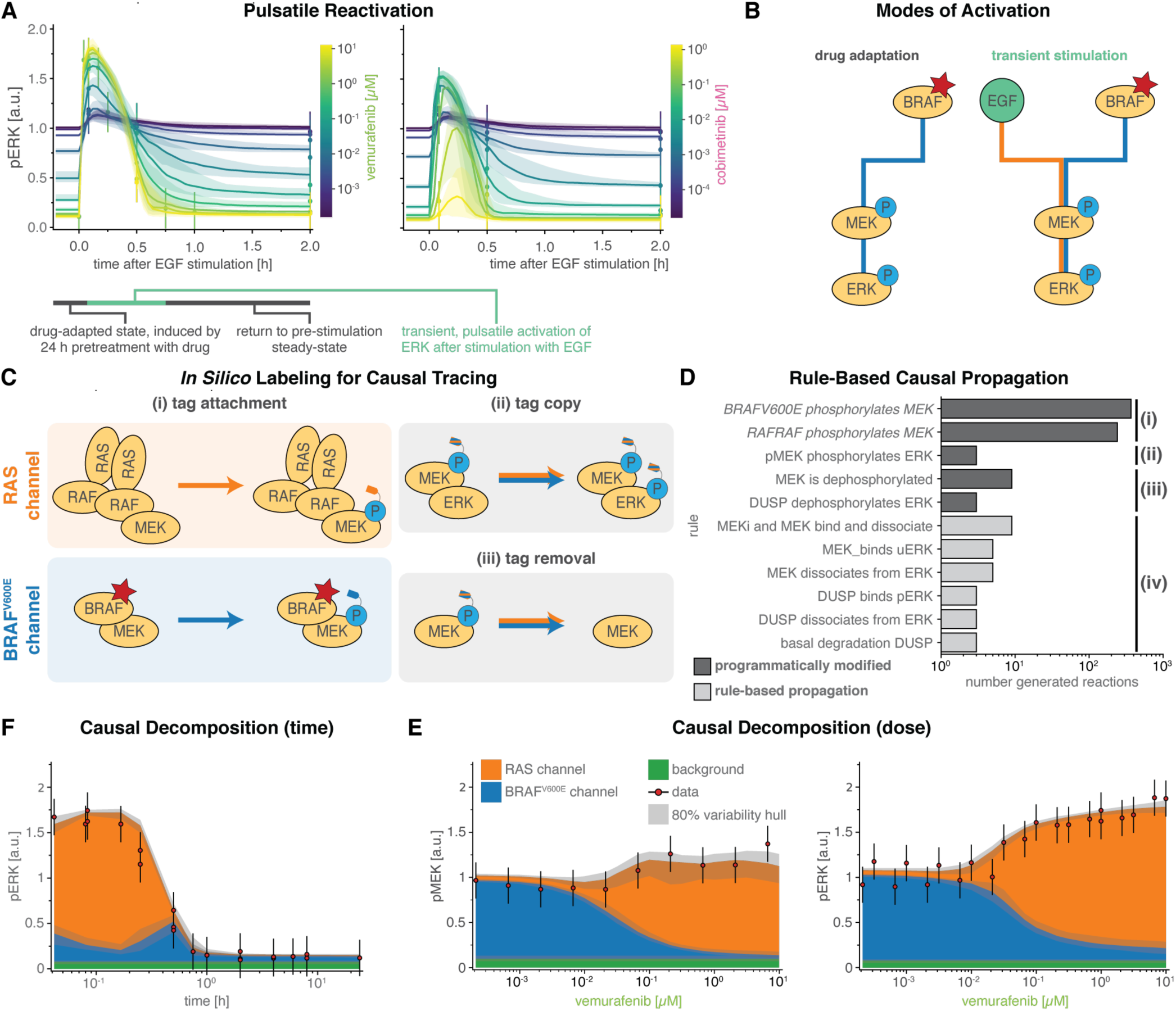
Causal Decomposition of RAS and BRAF^V600E^ Channels. **(A)** Time course of pre- and post- stimulation pERK levels. Model simulations are shown as solid lines, experimental data as vertical point- ranges. Colors indicate different concentrations of vemurafenib (RAFi) and cobimetinib (MEKi). Shading shows 80% percentiles over parameter sets. **(B)** Toggling of modes activation for pMEK via BRAF^V600E^ (blue) and EGF (orange) during the two phases of pulsatile reactivation shown in A: drug adaptation (left) and transient stimulation (right) **(C)** Schematic for tracing of causal history using synthetic sites. **(D)** Rules affected by causal decomposition **(E, F)** Comparison of experimental data and decomposed model simulations at 5 minutes after EGF stimulation. Data is shown as point-ranges. Median (over parameter sets) simulations are shown as stacked areas with color indicating reaction channel (blue: BRAF^V600E^, orange: RAS). Shading indicates 80% percentiles over parameter sets.

### Causal Decomposition untangles Intertwined BRAF^V600E^ and RAS Driven Signaling

When cells were adapted to RAFi (vemurafenib unless otherwise noted) for 24 hours, steady-state pERK levels decreased with drug concentrations. In striking contrast, the amplitude of pERK pulses generated by adding exogenous EGF increased with RAFi concentration (**Figure 2A** left). Thus, EGF (and other growth factors applied in a similar manner) induced pERK in proportion to the degree of BRAF^V600E^ inhibition. When MEKi (cobimetinib unless otherwise noted) was used over a dose range, a biphasic response was observed: below ∼0.1 µM MEKi EGF-induced pERK levels increased with MEKi concentration but above ∼0.1 µM MEKi they fell (**Figure 2A** right). In all cases, the effects of EGF were transient and pERK levels returned to their drug-adapted baseline levels within one to two hours. The calibrated MARM2.0 model recapitulated all of these phenomena and we therefore sought a molecular explanation using model analysis.

Experimentally determined pMEK and pERK levels measure the sum of active MAPK kinases generated by oncogenic and chronically active BRAF^V600E^ and by transiently active EGFR (**Figure 2B**). To decompose these two sources of MAPK activity, we modeled a “*RAS reaction channel*,” which encompasses all reactions initiated by (RAS-GTP)_2_-RAF_2_ oligomers, and a “*BRAF^V600E^ reaction channel*” encompassing all MAPK reactions downstream of the BRAF oncogene. In agent-based modeling, it is straightforward to keep track of the different origins of a single molecular species and thereby generate causal traces or “stories” (Boutillier *et al*, 2018). To adapt this approach to an ODE model, we used an *in silico* labeling strategy that involved adding a virtual “tag” to pMEK (**Figure 2C,** Methods Section Causal Signal Decomposition) at the time of its generation by (RAS-GTP)_2_-RAF_2_ (orange, top left panel) or BRAF^V600E^ (blue, bottom left panel). The tag was copied from pMEK to pERK upon ERK activation (blue/orange, top right panel) and removed during dephosphorylation (blue/orange, bottom right panel). Implementing this approach required modification of only of a few PySB rules (**Figure 2D**) and did not change model dynamics.

For causal decomposition of MARM2.0 under a range of conditions, computational labeling of both pMEK and pERK was necessary, since the two active forms do not have the same proportionality (degree of amplification) in the two reaction channels: in the BRAF^V600E^ channel, the MEK phosphorylation rate is lower when MEKi is bound to uMEK, generating a lower ratio of pMEK-MEKi to *apo*-pMEK than in the RAS channel, in which the MEK phosphorylation rate is independent of MEKi binding. The origins of this phenomenon are described in greater detail below. Since MEKi inhibits the catalytic activity of pMEK, amplification from pMEK to pERK is lower in the BRAF^V600E^ than the RAS channel.

The value of causal decomposition was illustrated when we investigated the observed increase in pERK levels in the BRAF^V600E^ channel following EGF addition (blue, **Figure 2E**). This was unexpected, since, in MARM2.0, EGF only activates the RAS channel. We surmised that activation of the BRAF^V600E^ channel might arise from retroactivity (Del Vecchio *et al*, 2008), in which downstream reactions affect upstream or parallel reactions by imposing a load on them, most commonly by competing for a limited pool of a regulators (Sauro, 2008). Using a counterfactual model, we confirmed that retroactivity in the BRAF^V600E^ channel arose from sequestration of DUSP proteins by pERK in the RAS channel (**Figure S2E**). Thus, activation of the RAS channel can activate the BRAF^V600E^ channel by reducing the rate of DUSP- dependent pERK dephosphorylation. A second example of causal decomposition involved experimental data showing that pMEK levels remain roughly constant over a 10^5^-fold range of RAFi concentrations (as monitored at the 5-minute peak of an EGF-induced pulse, **Figure 2F left**). Causal decomposition showed that this unexpected behavior arose from a steady reduction in the activity of the BRAF^V600E^ channel (blue) with increasing RAFi and a simultaneous and offsetting increase in signaling in the RAS channel (orange). This was true of all 3 RAFi and 5 MEKi tested (**Figure S3**) and represents a classic case of pathway rewiring that is obscured at the level of total MAPK activity.

### Slow Transcriptional Feedbacks Imprint Drug-Adapted State and Unravel Cyclic Causal Dependencies

Experimental data (Gerosa *et al*, 2020; Lito *et al*, 2012; Pratilas *et al*, 2009) and model trajectories show that DUSP (blue), SPRY (orange), and EGFR (green) proteins (dark colors) and mRNA (light colors) levels are substantially lower in cells adapted to RAFi for 24 hours as compared to drug-naïve cells (**Figure 3A left, S2B,F**). This is consistent with the known role of MAPK activity in promoting the expression of negative (feedback) regulators. However, it raises the question: why is pERK only transiently activated by EGF in drug-adapted cells if feedback is suppressed? When we simulated the induction of ERK pulses by exogenous EGF in drug adapted cells, we observed modest increases in EGFR, DUSP and SPRY mRNA levels (**Figure 3A right**), consistent with respective experimental training data (**Figure S2F**). However, at the protein level DUSP and SPRY remained almost constant and EGFR decreased. We surmised that this reflected the operation of transcriptional feedback on a longer time-scale (>2 h) than a typical EGF-mediated pulse (30-90 min). Model analysis showed that changes in EGFR protein levels were a consequence of receptor endocytosis, and degradation. Thus, EGFR trafficking and not negative feedback controls the duration of a pERK pulse in drug adapted cells, consistent with existing models of EGFR (Starbuck & Lauffenburger, 1992; Dessauges *et al*, 2021) and other transmembrane receptors (Becker *et al*, 2010). However, on the longer time-scale of drug adaptation, transcriptional feedback is the primary determinant of pERK levels. Similar separations in time-scale have been previously observed in other aspects of EGFR and MAPK signaling. For example, individual kinase phosho-states turn over on time scale of seconds but measurable changes in MAPK activity are a least hundred-fold slower, requiring minutes to hours (Kholodenko *et al*, 1999; Reddy *et al*, 2016; Kleiman *et al*, 2011). Thus, slow population average responses mask underlying biochemical reactions happening on much faster timescales.

**Figure 3:**
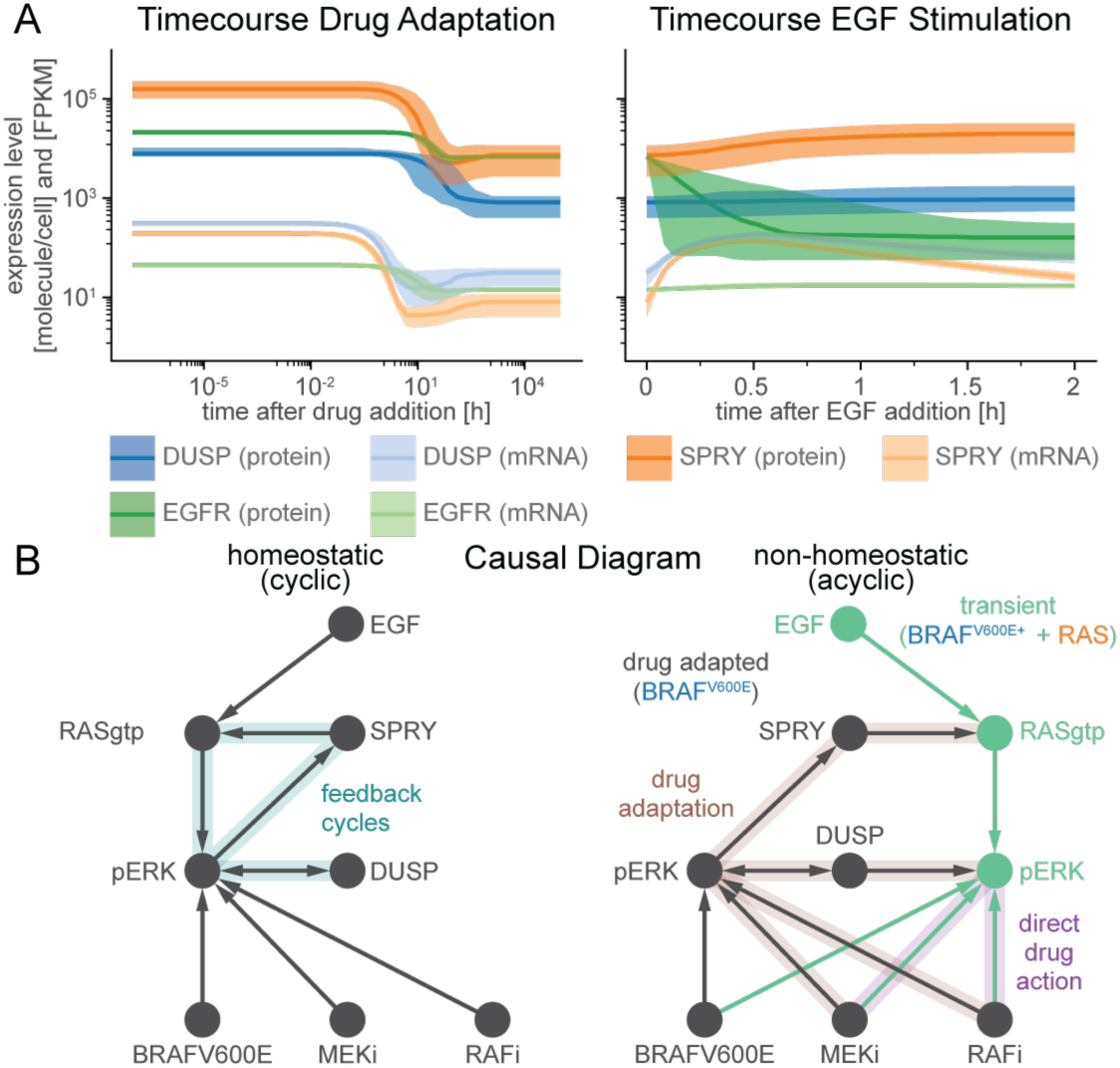
Transcriptional feedbacks imprint a sparse drug-adapted state. **(A)** Time Courses of pre- (left) and post-stimulation (right) protein (dark colors) and mRNA (light colors) expression levels of genes that are subject to transcriptional control by pERK. **(B)** Schematic of the structural causal model for the effect of RAFi and MEKi on pERK under homeostatic (left) and non-homeostatic (right) conditions.

The presence of feedback loops in a network usually generates cycles in the causal diagram (Mooij *et al*, 2013) (**Figure 3B** left), complicating model analysis (Pearl & Dechter, 2013; Spirtes, 2013). In the case of MARM2.0, a cycle involving positive regulation of feedback regulators by MAPK activities means, for example, that pERK activity could ultimately control DUSP levels or DUSP levels could control pERK activity. However, time-scale separation makes it possible to generate an acyclic causal diagram for MARM2.0 (Hyttinen *et al*, 2012) (**Figure 3B** right), in which the effects of RAFi and MEKi on pERK are split into the rapid and immediate effects of drug on kinase activity (*direct drug action*, purple shading) and a slower process involving changes in the levels of feedback proteins (*drug adaption*, brown shading). Prior to EGF stimulation, when only the BRAF^V600E^ channel is active (**Figure 2B** left), MEKi and/or RAFi levels control pERK levels in drug-adapted cells (*drug adapted pERK*; gray in **Figure 3B**), which in turn determine DUSP and SPRY concentration and, thus, the strength of negative feedback on pERK in the RAS channel (*transient pERK*, turquoise in **Figure 3B**). The indeterminacy between drug adapted pERK and DUSP levels remains (illustrated by a bidirectional edge in the graph), but this does not affect the determinacy between drug-adapted DUSP and transient pERK levels. Thus, time scale separation during drug adaption makes it possible to control the MAPK module in two distinct ways depending on the activating signal.

### MAPK Signaling is rewired by Drug Adaptation and Direct Inhibition

The ratio of input to output signals in a network (the gain) is a fundamental property of a signal transduction system that can be used quantify rewiring. Gain often varies along a series of reactions in a single channel – for example the number molecules of pERK generated per molecule of RAS-GTP as compared to EGF ligand. Gain could in principle be quantified by sensitivity (Goldbeter & Koshland, 1981), but as a mathematical concept, sensitivity is defined at steady-state, whereas signaling in the RAS channel is transient. Sensitivity could also be computed pointwise at every time point (Chen *et al*, 2009), but this would not account for the fact that input and output signals for any specific step in a network often have different timescales. For example, modeling revealed conditions in which an input signal (e.g., pEGFR levels) had started to fall following EGF stimulation, while a downstream event (e.g., formation of active RAS-GTP) was still increasing. We therefore defined the gain of a reaction channel as the ratio of L_∞_ or L_1_ norms (with respect to a logarithmic timescale) between input and output signals in corresponding model trajectories (see Methods; Signaling Gain). The L_1_ norm quantifies the area under the curve of the signal whereas the L_∞_ norm quantifies the height of the peak of the signal. Both represent scalar, time-independent quantities. For simplicity, we normalized gain to equal 1 in the absence of inhibition.

Gain for each of the two MAPK reaction channels can be investigated graphically using a formalism in which each node represents a “signal” that is defined as the sum of active model species, and edges represent signaling steps that are defined as the action of one or more PySB reaction rules. Gain was computed along each edge of the graph by computing the ratio of norms of input and output nodes. The graph in **Figure 4A** has been arranged so that each signaling step (edge) is affected by as few drug actions as possible – ideally only one - allowing changes in gain to be attributed to direct drug action (purple) or drug adaptation (brown). The graph contains three steps for the RAS channel (orange; steps R1-R3) and two steps for the BRAF^V600E^ channel (blue; steps B2-B3) with the channels “aligned” at the third step (pMEK phosphorylation of ERK; **Figure 4A**). We then used the calibrated model to compute time-resolved signals for all nodes at multiple drug concentrations (**Figure 4B**) and determined the gain (**Figure 4C)**. To visually summarize the inhibitor and concentration-dependent states of the graph, we generated separate representations for RAFi (**Figure 4D)** and MEKi (**Figure 4E**), with signal activity indicated as node opacity and gain as edge opacity.

**Figure 4:**
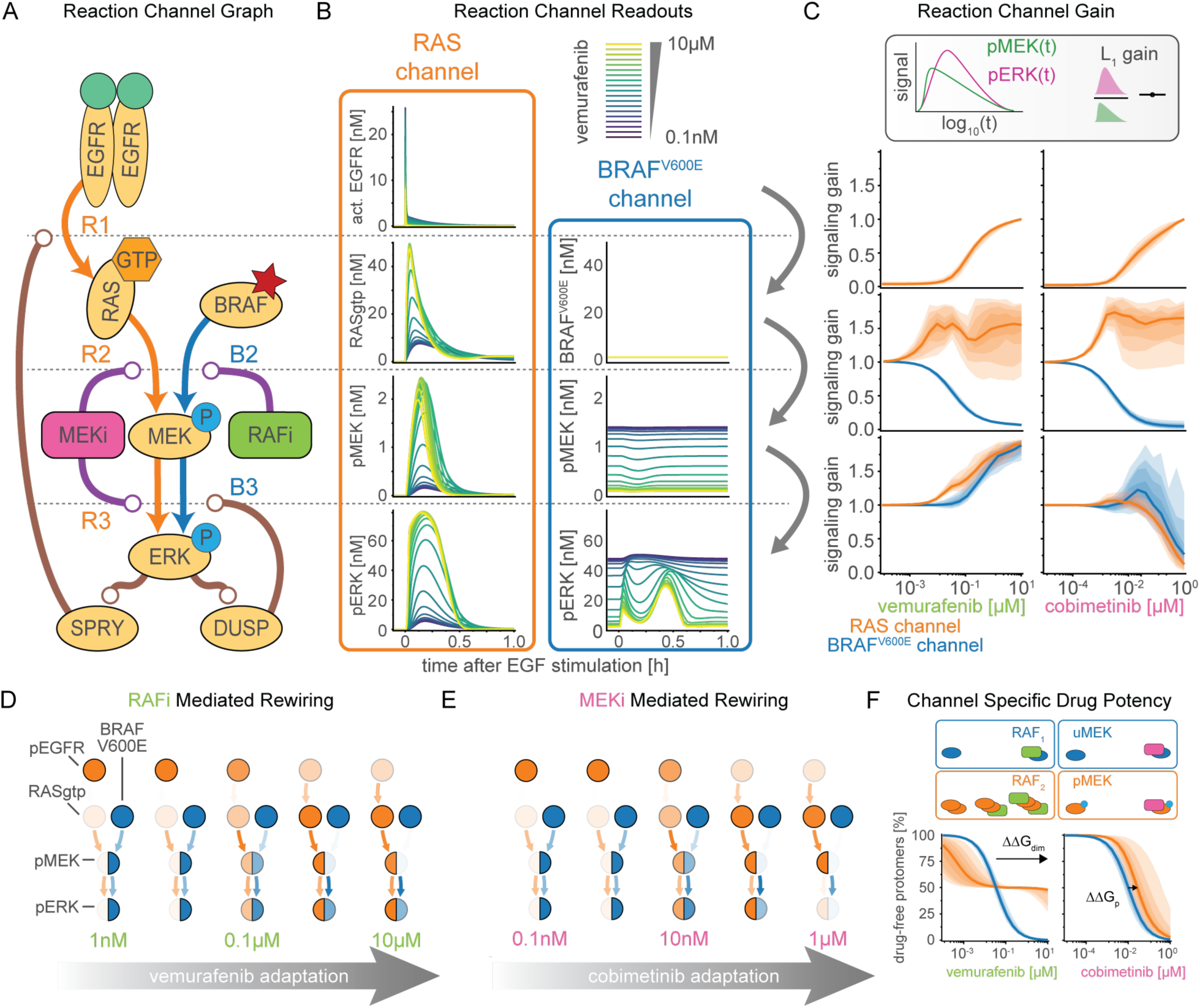
Quantification of signal transduction in RAS and BRAF^V600E^ channels. **(A)** Simplified model network depicting intertwined RAS and BRAF^V600E^ channels and feedbacks. **(B)** Decomposition of RAS and BRAF^V600E^ signals at the different nodes of the simplified network from A for different concentrations of vemurafenib. Color indicates vemurafenib concentration. Simulations were performed for a representative parameter value. **(C)** Quantification of signal transmissions in terms of signaling gain along the edges of the simplified network in A for different concentrations of vemurafenib (left) and cobimetinib (right). Color indicates the reaction channel. Shading indicates 20%, 40% 60% and 80% percentiles over parameter sets. **(D, E)** Visualization of pathway rewiring as a result of drug adaptation. Opacity of nodes indicates median normalized signaling activity (shown in B). Opacity of arrows indicates median normalized signaling gain (shown in C) where 100% corresponds to a signaling gain of 2. **(F)** Quantification of efficacy of drug inhibition. For RAF dimers, each protomer is counted individually. Shading indicates 20%, 40% 60% and 80% percentiles over parameter sets.

We found that drug adaptation to RAFi and MEKi had a similar impact on the first step (R1) of both reaction channels (**Figure 4C,** top panels). At low to medium drug concentrations (RAFi: ∼10^-4^ to10^-2^ μM, MEKi ∼10^-5^ to 10^-3^μM), the gain from pEGFR to RAS-GTP was close to zero representing complete inhibition of EGF-mediated signaling by the combined actions of feedback regulators such as DUSP and SPRY. At medium to high drug concentrations (RAFi: ∼10^-2^ to 10^-1^μM, MEKi: ∼10^-3^ to 10^-0^μM) a reduction in the levels of feedback regulators led to a relief of feedback and an increase in gain. At the second step, for medium to high RAFi and MEKi concentrations, we found that B2 had gain close to zero, but R2 gain was larger than one (**Figure 4C,** middle panels), indicating channel-specific effects for both drugs. For RAFi, we attributed this channel specificity to difference in the affinity of the RAFi for monomeric RAF in the BRAF^V600E^ channel and dimeric RAF in the RAS channel (orange vs. blue colored nodes). The difference in affinity is determined by the thermodynamic parameter ΔΔ*G*_234_ (**Box 3**), which encodes the ratio of drug affinities for the first and second protomers of a RAF dimer; for vemurafenib this difference was estimated to be ∼2.5 x 10^3^-fold (median of values from best 5% of fits). Thus, even at 10μM, the highest vemurafenib concentration tested, and a value well above the clinically useful range, ∼25% of RAF dimers had one protomer not bound to drug (**Figure 4F**, left), a configuration that is active as a kinase (Karoulia *et al*, 2017). The estimated lower bound for ΔΔ*G_dim_* corresponding to ∼60 fold decrease in affinity is consistent with a previously reported values of 30-100 fold lower IC_50_ for a BRAF^V600E^ relative to wild-type, as estimated from cell-based experiments with a splicing-variant that forms BRAF^V600E^-BRAF^V600E^ dimers (Karoulia *et al*, 2016). Moreover, estimated ranges for ΔΔ*G_dim_* were similar for the four other type I**½** RAFi drugs we tested (**Figure S1, S4**). For MEKi, we attributed the channel specific potency in the second step to a decrease in MEK phosphorylation rate by BRAF^V600E^ for BRAF-uMEK-MEKi complexes as compared to BRAF-uMEK complexes; modeling suggested a ∼ 6.5 x 10^3^-fold reduction in rate of reduction as compared to apo MEK with cobimetinib as the MEKi. Estimated values were similar (>800 fold) for trametinib and PD0325901, but substantially lower (<200 fold) for binimetinib and selumetinib, consistent with previously reported differences in the activity of these drugs (Pino *et al*, 2021). In all cases, the combination of lower RAFi affinity or lower MEKi-depdendent phosphorylations rate resulted in incomplete inhibition of pMEK in the RAS channel (**Figure S3**).

For the third step, we found that gain from pMEK to pERK (B3 and R3) increased at medium to high concentrations of RAFi (**Figure 4C,** bottom left panel), due to a reduction in DUSP expression levels. In contrast, MEKi did not have any effect on gain at medium concentrations (∼10^-3^ to 10^-2^μM, (**Figure 4C,** bottom right panel). This was unexpected, since the analysis described above shows that DUSP levels are controlled by drug-adapted pERK levels, which are inhibited at medium concentrations of MEKi and RAFi (blue, middle panels, **Figure 4C**). However, B3/R3 are the only steps in which the model implements two distinct effects for each drug: increases in ERK activity as a result of drug adaptation, i.e., DUSP downregulation, (brown, **Figure 4A**) and reductions in ERK activity via direct drug action by MEKi on MEK (purple, **Figure 4A**). Modeling suggested that direct drug action and adaptation balanced each other at intermediate MEKi concentrations and direct inhibition became dominant only at high concentrations. The differential potency of MEKi for BRAF^V600E^ (B3, blue) compared to the RAS channels (R3, orange) could also be due to a difference in affinity of MEKi for pMEK as compared to uMEK (see Section on Causal Decomposition) (Hatzivassiliou *et al*, 2013), which was encoded in the thermodynamic parameter ΔΔ*G*_5_ (**Figure 4F**). For cobimetinib, the inferred ΔΔ*G_p_* values corresponded to a ∼3.5-fold decrease in affinity, but for the four other MEKi tested this difference was >10-fold (**Figure S4**). Since the shift in MEKi potency for pERK activated by EGFR as compared to BRAF^V600E^ activated pERK was ∼100 fold (**Figure 2A, S2C**), we concluded that it likely arises from a combination of channel specific efficacy in the second step and balancing of direct drug action and drug adaptation in the third step. In this form of the model, the decreased affinity of MEKi for pMEK played only a minor role in channel specificity.

One interesting aspect of gain in MAPK signaling is that it varied independent of total activity of the signaling cascade or the flux of MAPK kinases and phosphatases (**Figure 4D,E**). For example, at high concentrations of RAFi, step B3 had high gain (due to low DUSP activity) but the channel was functionally inactive (due to RAFi-BRAF^V600E^ binding). The interesting feature of this arrangement is that the anti-proliferative effects of RAFi are highly sensitive to anything able to activate MEK directly, such as a mutation in the kinase. Consistent with this, activating mutations such as MEK1^C121S^ are observed to give rise to acquired drug resistance in patients (Wagle *et al*, 2011). High gain but low activity in the RAS channel is directly analogous, and potentiates both ligand-mediated RTK activation and RAS mutation (e.g., NRAS^Q61K^ discussed below). More generally, it is possible that identifying signaling steps with low activity but high gain may help to pinpoint mechanisms of potential acquired drug resistance.

### Pulsatile Signaling Induces Apparent Drug Interactions

MEK and RAF inhibitors are normally used in combination. To study drug interaction and also test the predictive power of MARM2.0 in conditions distinct from those used for model training, we simulated the effects of RAFi plus MEKi combinations on pERK levels with a model trained on single-drug responses alone (the model training described above). Drug dose-response relationships were then visualized as surface plots (**Figure 5A**) and isobolograms (**Figure 5B**). In the absence of exogenous growth factors (**Figure 5A(i)**), we predicted a monotonic decrease in pERK levels with increasing doses of both drugs (left panels) and experimental data were in agreement (right panels). In BRAFi- adapted and EGF stimulated cells, we predicted a more complex landscape (**Figure 5A(ii)**), in which pERK was relatively drug resistant along a L-shaped region (red dashed outline) at intermediate MEKi and high RAFi concentrations with a gradual decrease at high MEKi concentrations. Using isobolograms, we observed disconnected level sets (bottom, **Figure 5B**), recapitulating the non-monotonic response to MEKi in **Figure 2A,** in which pERK levels first rose and then fell with increasing drug concentration. Experimental data (right panel, **Figure 5A(ii)**) was qualitatively similar to predictions (left panel) and differences were primarily in the magnitude of pERK, not the shape of the response surface (bottom, **Figure 5B**). Disconnected isobolograms (bottom, **Figure 5B**) are noteworthy, because measures of drug interactions such as Loewe additivity (Loewe, 1928) or the Chou-Talalay combination index (Chou *et al*, 1993) require a one-to-one mapping between dose and response (a bijective curve) and cannot be applied in this context. However, comparing pERK levels to null models for Bliss independence (Bliss, 1939) (Bliss, **Figure 5C**) and highest single agent (Lehár *et al*, 2007) (HSA, **Figure 5D**) revealed negligible drug interaction (white) in the absence of EGF (top panels) in simulation (left) and experimental data (right). Under conditions of EGF stimulation (bottom panels), we observed substantial discordance between the magnitude and sign of drug interaction as scored by Bliss criteria (**Figure 5C**) and HSA (**Figure 5D**). Thus, existing definitions of drug synergy and antagonism do not adequately describe the complex dose-response landscapes we observed.

**Figure 5:**
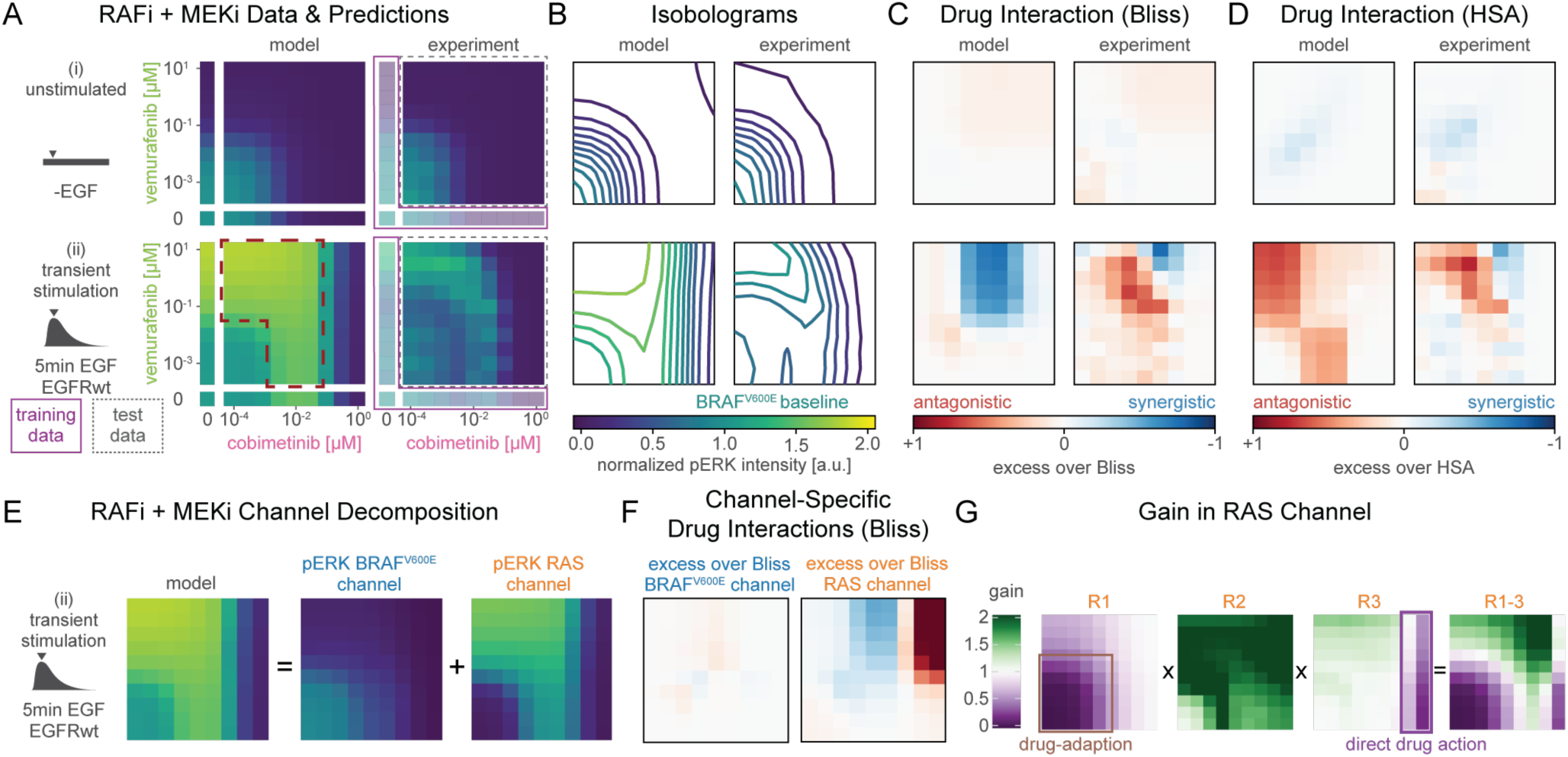
Prediction and analysis of drug combinations. **(A)** Experimental data and model simulations for ERK combination response without EGF stimulation (top) and 5 min after EGF stimulation (bottom). Training data has lower opacity and purple outline. Test data has a grey, dashed outline. **(B)** Isobolograms of smoothed dose response surfaces from A. Concentrations and color scheme are the same as in A **(C, D)** Analysis of drug synergy according to excess over Bliss and highest single agent (HSA). Concentrations are the same as in A. **(E)** Decomposition of pERK model simulations at 5 min after EGF stimulation (left) in BRAF^V600E^ (middle) and RAS (right) channels. Color and concentrations are the same as in A. **(F)** Drug interaction analysis for decomposed channels. Color is and concentrations are the same as in A. **(G)** Quantification of signaling gain in the physiological signaling challenge. Pointwise multiplication is indicated by *x*. Reaction steps (Figure 4A) are indicated on top. Purple and brown outlines indicate molecular mechanisms responsible for lower gain. Concentrations are the same as in A.

When we decomposed dose-response surfaces for EGF-stimulated conditions (left, **Figure 5E**) into BRAF^V600E^ (middle) and RAS channels (right). we observed little RAFi and MEKi interaction in the BRAF^V600E^ channel (left, **Figure 5F**) and either a small level of synergy (blue) or strong antagonism (red) in the RAS channel depending on drug concentration (right). When we then computed gain in the RAS channel for R1, R2 and R3 (**Figure 4A**) at different drug concentrations, we observed low gain for R1 at RAFi and MEKi concentrations below 10 and 1 nM respectively (first panel, **Figure 5G**), high gain for R2 at all concentrations (second panel) and low gain for R3 at MEKi at >1μM (third panel). When the gain for steps R1-R3 was computed as pointwise multiplication of the three surfaces, the L-shaped region of drug resistant pERK (fourth panel) was regenerated (**Figure 5A(ii)**). Thus, the overall drug response landscape can be explained by the superposition of adaptive drug response on R1 (brown), and direct drug effects on R3 (purple).

### Sustained Signaling does not Induce Drug Interaction

To study the effects of RAFi and MEKi on signaling in the RAS channel under conditions of sustained rather than transient EGFR activation, we over-expressed EGFR using CRISPRa (Gerosa *et al*, 2020), yielding two cell lines with 4-fold (light blue) and 9-fold (turquoise, referred to as A375 CRISPRa-EGFR below) increases in expression levels (**Figure 6A**). It has previously been shown that, when EGFR is overexpressed to this degree, mechanisms of receptor endocytosis and degradation are saturated and EGFR becomes chronically rather than transiently active in the presence of ligand (Lund *et al*, 1990; Wiley, 1988; Kiyatkin *et al*, 2020). Consistent with this, we found that upon ligand addition, pERK levels in RAFi-adapted CRISPRa-EGFR cells rose rapidly to a peak at ∼30 min and then fell slightly to level at roughly ∼75% of their levels in the absence of RAFi exposure; pERK remained at this level for at least 24h in both experiments and simulations. Under these conditions, RAFi had substantially lower efficacy (EC_max_; **Figure 6B**) and MEKi had lower potency (EC_50_; **Figure 6C**) than in cells not stimulated with EGF. Channel decomposition (**Figure 6B** right panels) revealed an increase in pMEK and pERK levels in the RAS channel (orange) and also in the BRAF channel (blue), which we ascribed to retroactivity (**Figure S6C**) and low DUSP levels (**Figure 6A**, bottom, dark blue). Analysis of pERK phase space with DUSP and SPRY mRNA levels and SPRY protein levels showed similar distributions at 8h post EGF-stimulation in drug-adapted CRISPRa-EGFR cells and pre EGF-stimulation in drug-adapted EGFR^wt^ cells, suggesting a steady state had been reached at 8h post EGF-stimulation (**Figure S6D**). In contrast, DUSP protein levels at 8h post EGF-stimulation remained up to 3-fold below the levels observed at the same pERK levels pre EGF-stimulation, suggesting steady-state had not yet been reached, which is consistent with long DUSP protein half-life times observed in Western blot experiments (Lito *et al*, 2012). Thus, the relative resistance of EGFR amplified cells to RAFi and MEKi appears to result from sustained activation of the RAS channel and slow DUSP protein turnover.

**Figure 6:**
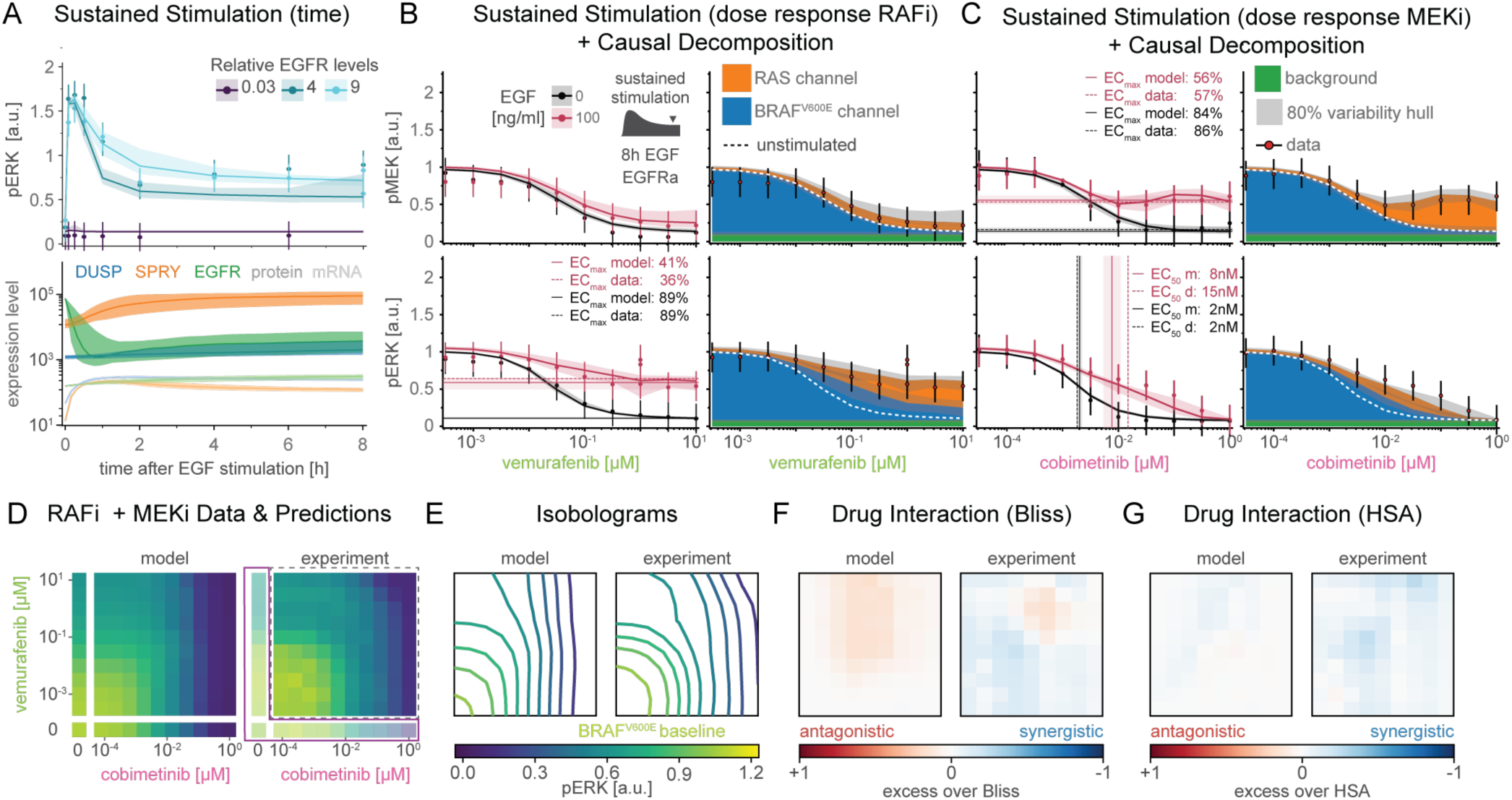
Prediction of resistance from EGFR upregulation. **(A)** Prediction of time course data for three different clones (two overexpression, one knockdown). Solid line show medians. Shading indicates variability across 80% of parameter sets. Top plot shows pERK response. Bottom plot shows mRNA (light colors) and protein (dark color) expression level changes. **(B, C)** Prediction of dose response data with and without EGF at 8 hours after stimulation in response to vemurafenib (B), cobimetinib (C). Left panels show EGF stimulated (red) and unstimulated (black) conditions. Right panels show decomposed model simulations for EGF stimulated conditions. Data is shown as point-ranges. Median (over parameter sets) simulations are shown as stacked areas. Shading indicates 80% percentiles over parameter sets. Simulations for EGF unstimulated conditions are shown as white dashed line. **(D)** Experimental data and model simulations for pERK combination response at 8h after EGF stimulation. Training data has lower opacity and purple outline. Test data has a grey, dashed outline. **(E)** Isobolograms of smoothed dose response surfaces from A. **(F, G)** Analysis of drug synergy according to excess over Bliss (F) and HSA (G).

When we predicted the pERK dose-response surface for combined RAFi and MEKi treatment of CRISPRa-EGFR cells (8h after stimulation with EGF) using single drug training data (**Figure 6D** left), we observed incomplete pERK inhibition at high RAFi and medium MEKi concentrations. The resulting isobolograms had a convex shape (**Figure 6E**) with minimal drug interaction by Bliss (**Figure 6F**) or HSA criteria (**Figure 6G**). This differs from what was observed with pulsatile RTK activation (**Figure 5C, D** bottom panels) and suggests that drug interactions in the case of pulsatile signaling were only possible due to time scale separation between drug adaption and direct drug action.

### Structure-Based Model Formulation Enables Generalization Across Inhibitor Classes

In MARM2.0, the thermodynamic parameter ΔΔ*G_dim_* describes changes in the stability of (RAFi-RAF)_2_ complexes; these have been studied in detail via crystallographic structures (Rukhlenko *et al*, 2018). Negative ΔΔ*G_dim_* values manifest themselves as a loss of drug affinity by the second protomer in a RAF dimer. It is well-established that this leads to lower RAFi efficacy in the RAS channel as compared to the BRAF^V600E^ channel (**Figure 4C,F**). However, due to energy conservation (**Box 3**), ΔΔ*G_dim_* <0 also results in a higher dissociation rate of RAF_2_ complexes at high RAFi concentrations (**Figure S5**). Thus, thermodynamically formulated models can describe the phenotypic response to inhibitors based on their allosteric properties. In contrast to type I**½** RAF inhibitors, type II inhibitors (also called panRAFi; **Box 2**) such as LY3009120 and AZ-628 (Henry *et al*, 2015; Noeparast *et al*, 2018) inhibit both monomeric RAF in the BRAF^V600E^ and dimeric RAF in the RAS channel with similar affinity. Crystallographic data suggest that this arises because panRAF inhibitors do not destabilize (RAFi-RAF)_2_ complexes, i.e., they do not induce allosteric changes. To determine whether MARM2.0 correctly predicts the response to type II inhibitors based on the loss of allostery, we calibrated MARM2.0 using data from A375 CRISPRa-EGFR cells that were treated with LY3009120 (**Figure 7A**) or AZ-628 for 24h (**Figure S6A**), but not stimulated with EGF (**Figure S2C**). This allowed estimation of drug affinity for monomeric RAF (Δ*G*); ΔΔ*G_dim_* was fixed to 0 to reflect loss of allostery. We then generated predictions for pMEK (top) and pERK (bottom) levels 8 hours after EGF stimulation (red) in cells adapted to LY3009120 (**Figure 7A** left panels) or AZ-628 (**Figure S7**). Predictions matched experimental data under the same conditions and causal decomposition confirmed that RAF was strongly inhibited in the RAS channel (right panels). We also observed good agreement between model predictions and experimental data for LY3009120 in combination with cobimetenib in EGF-stimulated, drug-adapted cells (**Figure 7B**). Analysis of drug interactions using HSA and Bliss criteria (**Figure 7C**) revealed a similar level of additivity (but little or no synergy) in model predictions and experimental data (note that the isoboles are curved not due to synergy but our use of logarithmic concentration axes). These data show that MARM2.0 can correctly predict the properties of different RAF inhibitors based on differences in their allosteric properties alone.

**Figure 7:**
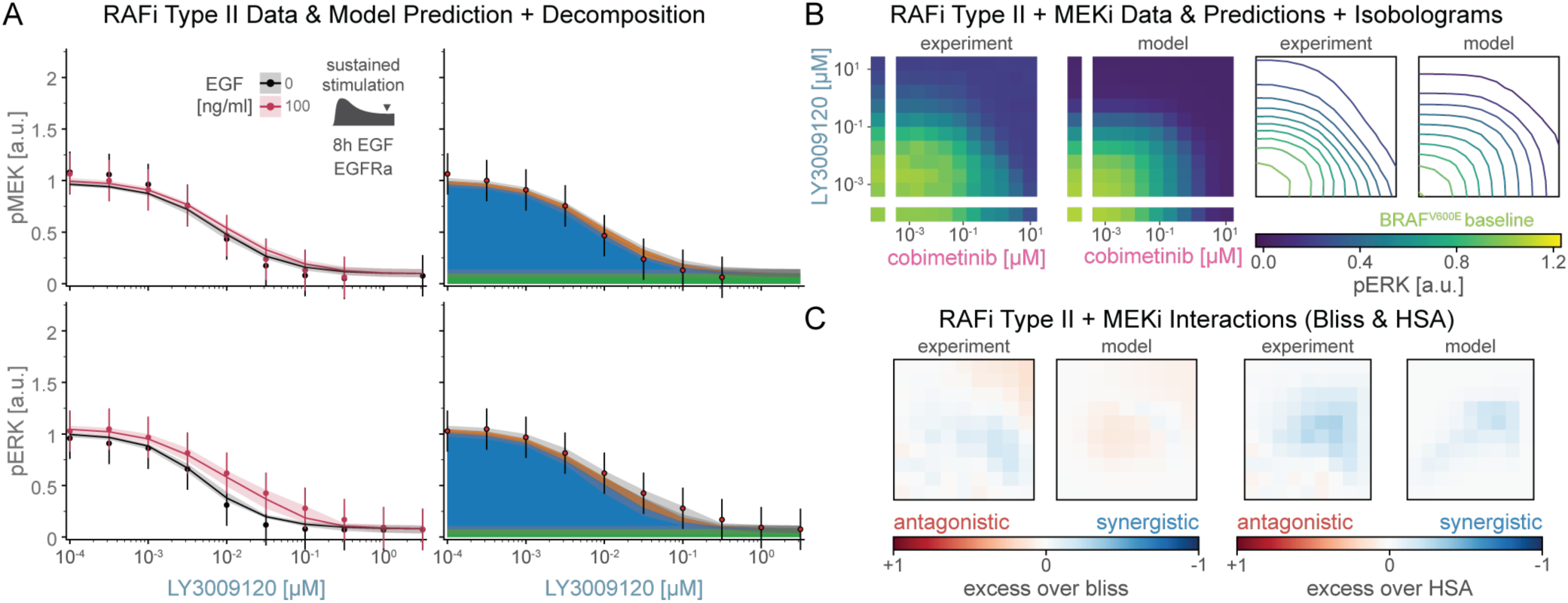
Prediction of response to panRAF inhibitor LY3009120. **(A)** Comparison of pMEK (top) and pERK (bottom) dose response predictions and experimental validation for A375 EGFR-CRISPRa with (red) and without (black) 8h of EGF stimulation. Solid lines and stacked areas show median (over parameter sets) simulations. Shading indicates 80% percentiles over parameter sets. Data is shown as point-ranges. **(B)** Drug combination response for A375 EGFR-CRISPRa 8h after EGF stimulation. **(C)** Analysis of drug synergy according to excess over Bliss (left) and HSA (right).

### Successes and Limitations in Extending MARM2.0 to Other Resistance Mechanisms

NRAS^Q61K^ is a frequently observed resistance mutations found in melanoma patients treated with RAF/MEK therapy (Long *et al*, 2014; Shi *et al*, 2014). We modelled NRAS^Q61K^ as RTK-independent activation of the RAS channel (Burd *et al*, 2014), with baseline pERK levels inferred from drug-naïve NRAS^Q61K^ BRAF^V600E^ double mutant melanoma cells (**Figure 8A**). Under these conditions, simulations recapitulated higher baseline pERK and predicted 7-fold lower efficacy for RAFi (NRAS^Q61K^, turquoise; left panels) and 4-fold lower potency for MEKi (**Figure 8B** right panels) as compared to NRAS wildtype cells (NRAS^wt^, purple). These predictions were confirmed in A375 cells engineered to conditionally express NRAS^Q61K^ (Yao *et al*, 2015), but the observed loss of MEKi potency was even greater than modeling predicted (30-fold). Causal decomposition of (modelled) pERK activity in the presence of drug combinations (varying MEKi plus 1μM RAF; **Figure 8C**) showed that 1μM RAFi was sufficient to completely block activity in the BRAF^V600E^ channel (blue) without affecting the RAS channel (**Figure 8B** and **8C**). This made it possible to study NRAS^Q61K^ signaling without interference from the BRAF^V600E^ oncogene.

**Figure 8:**
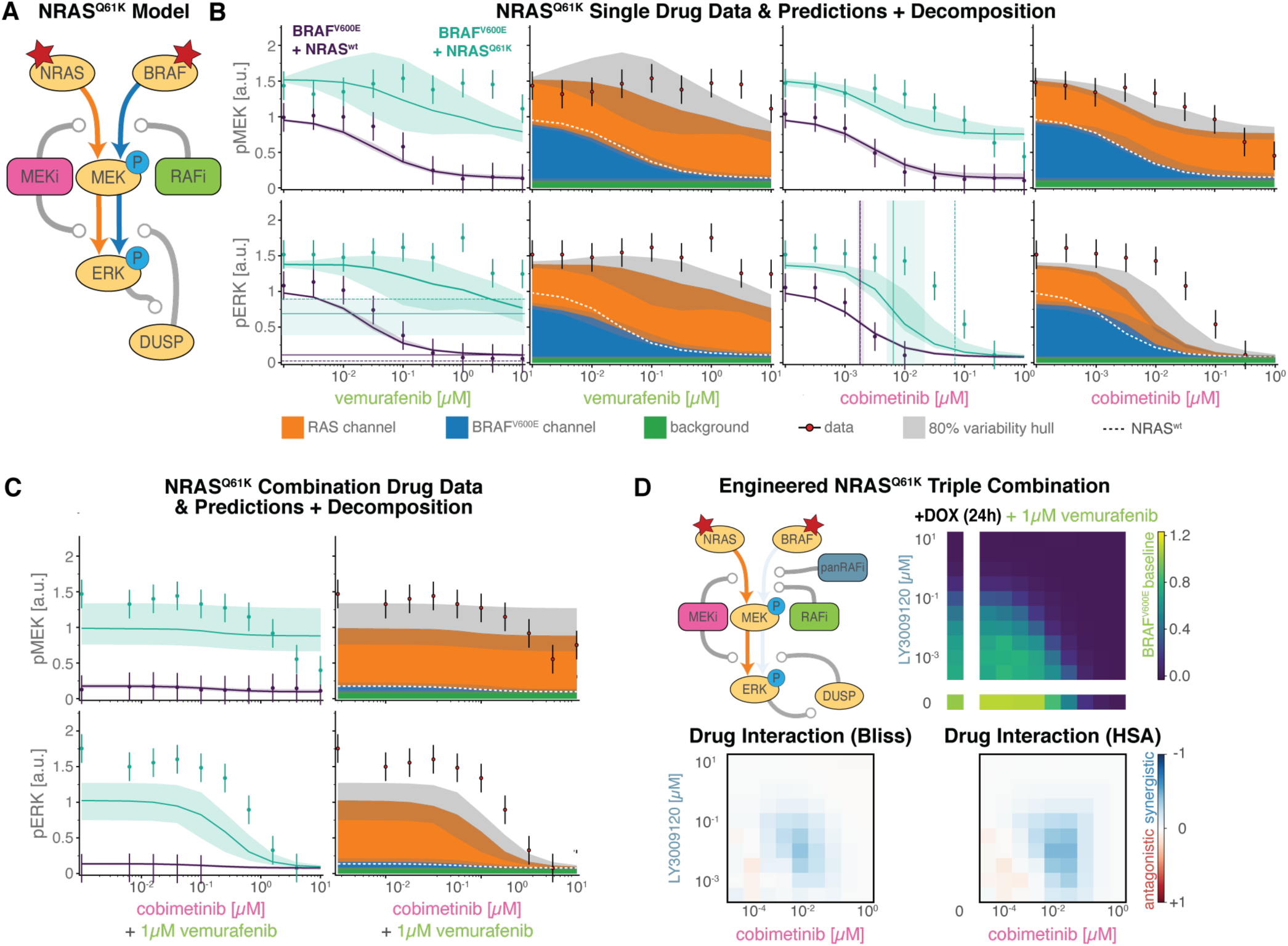
Prediction of response with NRAS^Q61K^ mutations. **(A)** Sketch of simplified model topology induced by NRAS^Q61K^ mutation. **(B, C)** Comparison of pMEK (top) and pERK (bottom) dose response predictions and experimental validation for A375 cells with inducible NRAS^Q61K^ mutation (induced: turquoise, uninduced: purple). Solid lines and stacked areas show median (over parameter sets) simulations. Shading indicates 80% percentiles over parameter sets. Data is shown as point-ranges. Vertical lines indicate EC_50_ values, Horizontal lines indicate EC_max_ values (data: dashed, model: solid). **(D)** Combination dose with 24h Dox stimulation to the triple combination of 1 μM RAFi (vemurafenib) plus varying doses of panRAFi (LY3009120) and MEKi (cobimetinib). Drug interaction analysis via Bliss (bottom left) and HSA (bottom right).

Based on this insight we devised a triple combination experiment to study drug interactions between panRAFi and MEKi in the RAS channel alone (**Figure 8D**, top left panel). A375- BRAF^V600E^ NRAS^Q61K^ cells were grown in the presence of 1μM vemurafenib plus different concentrations of LY3009120 and cobimetinib for 24h and pERK levels then determined (top right panel). In contrast to the analogous experiment without 1μM vemurafenib (**Figure 7C**), we observed pronounced synergy (blue) at low to medium concentrations of both inhibitors (∼1-100nM) by Bliss (bottom left panel) and HSA criteria (bottom right panel). Similar synergy has previously been observed in KRAS-driven cell lines of diverse origins (Yen *et al*, 2018). However, we found that the effects of combining three drugs in double mutant cells A375 cells were not accurately captured by MARM2.0 (**Figure S7**). We hypothesized that drug synergy is likely to arise due to a combined allosteric effect of both drugs on RAS-RAF-MEK complexes, as similar interactions have been described for combined treatment of MEKi and APS-2-79, a type II inhibitor of the KSR scaffolding protein (**Box 2**) (Dhawan *et al*, 2016). MARM2.0 does not include such allosteric effects and was not trained on combination data that would be necessary to infer the strength of the combined effect *a posteriori*. This limitation of MARM2.0 can be rectified in future studies, but serves to reveal how the subtleties of drug interactions can be relatively difficult to discern when multiple parallel reaction channels are active.

### Model for Melanoma Cell Line Generalizes to Colorectal Cell Line

BRAF^V600E^ mutations are found in a variety of cancers other than melanoma, notably colorectal cancers. To investigate whether MARM2.0 could predict the responses of BRAF^V600E^ colorectal cancers to RAFi, we collected data from HT29 cells, which carry a BRAF^V600E^ mutation and have high EGFR expression (similar to A375 EGFR-CRISPRa cells). We anticipated that BRAF^V600E^ channel would be a primary driver of pERK levels in the absence of EGF (**Figure 9A**) and the RAS channel in the presence of EGF (**Figure 9B**). To instantiate MARM2.0 for HT29 cells, we rescaled baseline protein and mRNA expression levels according to relative abundances in proteomic and transcriptomic data from the Cancer Cell Line Encyclopedia (Barretina *et al*, 2012; Nusinow *et al*, 2020). We simulated pERK drug response for RAFi plus MEKi combinations for HT29 cells (bottom) and compared to simulations for A375 CRISPRa-EGFR (top) and Dox inducible NRAS^Q61K^ A375 cells (middle). In all three cell lines, model predictions (left) demonstrated pERK inhibition in high-dose combinations, a result confirmed by experimental data (right; **Figure 9C**). Under conditions of EGF-stimulation, simulations and data revealed drug-resistant ERK activation (**Figure 9D**) and an ∼10-fold rightward shift in RAFi and MEKi dose-response curves (red arrows). Causal decomposition (**Figure 9E**) confirmed that these changes in drug potency are a consequence of profound differences between the BRAF^V600E^ (left) and RAS (right) reaction channels. The remarkably good agreement between MARM2.0 predictions and data in three different settings in which RAFi resistance is observed (EGF treatment in BRAF^V600E^ melanoma and colorectal cancer and NRAS^Q61K^ expression in BRAF^V600E^ melanoma) suggests that the model correctly unifies the key features of allosteric regulation of oncogenic MAPK signaling.

**Figure 9:**
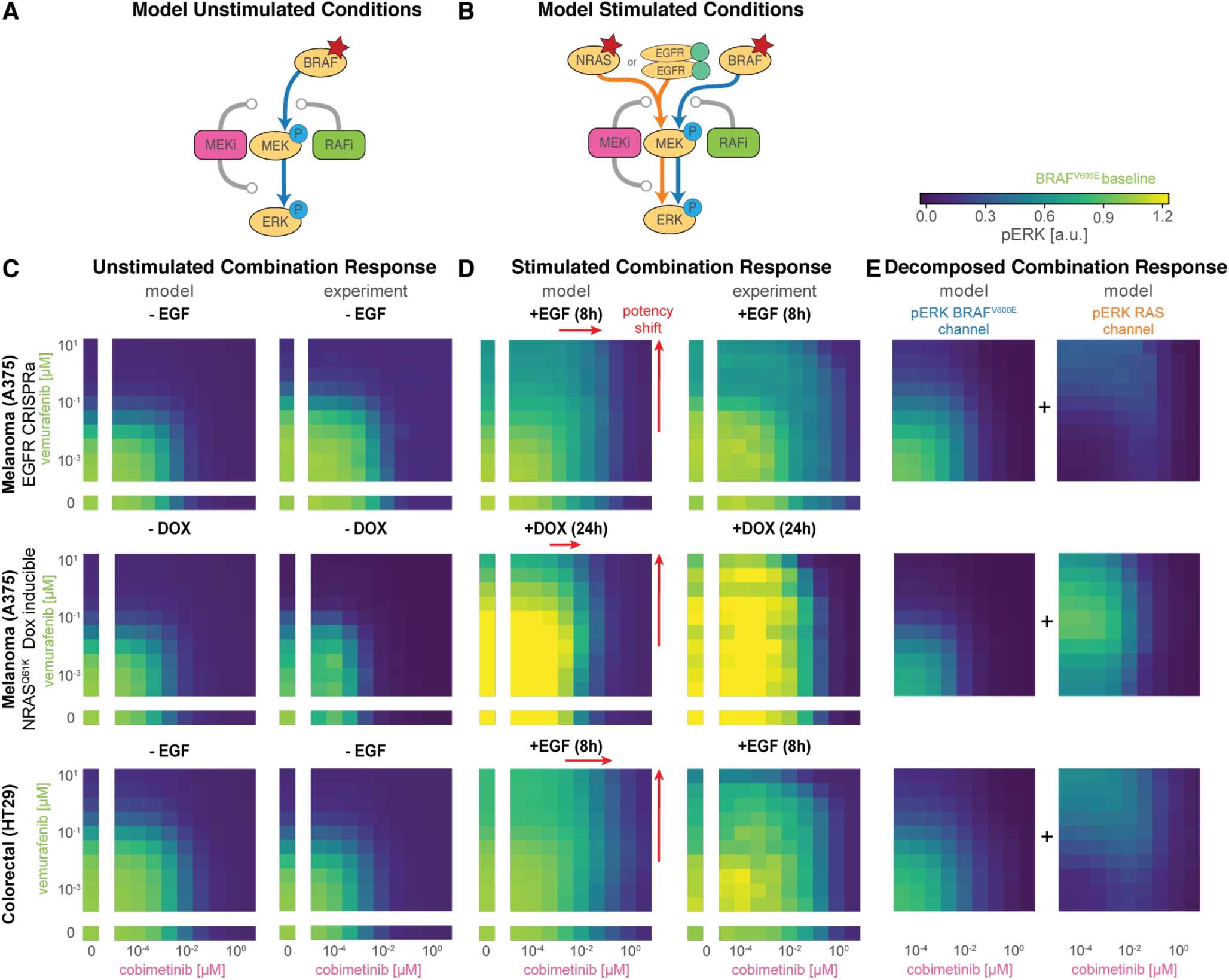
A unified model of drug resistance in BRAF-mutant cancers. pERK Drug combination response for (i) A375 melanoma cell line EGFR-CRISPRa amplified cell line with (right) or without (left) 8h of EGF stimulation (first row), (ii) A375 melanoma NRAS^Q61K^ Dox-inducible cell line with (right) or without (left) 24h Dox stimulation (second row) and (iii) HT29 colorectal cell line with (right) or without (left) 8h EGF stimulation (third row).

## DISCUSSION

In this manuscript we describe a quantitative framework for analyzing “pathway rewiring” with specific reference to rewiring involved in adaptive resistance to MEK and RAF inhibitors in BRAF^V600E^ melanoma. We described new analytical methods and a mass-action kinetic model (MARM2.0) that substantially extends previous models of MAPK signaling by using an energy-based formalism to efficiently represent allosteric regulation of MAPK kinases and the complexes they form with each other and with small molecule drugs. Among other analysis, we used MARM2.0 to predict and understand resistance in the context of an NRAS mutation that is frequently observed in melanoma patients who acquire resistance to MEK and RAF inhibitors and BRAF^V600E^ colorectal cancer cells that are intrinsically resistant to RAFi.

The MAPK cascade, and RTKs acting upstream of it, are among the signal transduction systems most intensively studied using systems of ODEs and dynamical systems analysis. Adaptive drug resistance in BRAF^V600E^ melanoma therefore represent an excellent setting in which to advance the state of the art in mechanistic modeling of intracellular networks. Moreover, whether adaptive or acquired, resistance to MEK and RAF inhibitors is directly relevant to patient outcomes: many individuals with BRAF^V600E^ melanoma experience rapid and live-saving remission with relatively little adverse effect. However, the frequent and rapid emergence of drug resistance (often within a year of the start of treatment) dramatically reduces survival. Preventing the acquisition of drug resistance is widely seen as the key to achieving more durable responses to MEK-RAF inhibitors and targeted anti-cancer drugs in general.

MARM2.0 fits well to over 900 data points in over 200 experimental conditions, requiring only eleven proteins, three mRNA species and three small molecule drugs. However, capturing the known activities, interactions, and structural features of these relatively few molecules involved a network of over 30,000 distinct biochemical reactions. MARM2.0 accurately predicted the responses of cells to ten different investigational and approved small-molecule kinase inhibitors in over 1600 experimental conditions, including drug combinations outside of the training dataset. While resistance in many of these conditions has been attributed to specific mechanisms (Haling *et al*, 2014; Hatzivassiliou *et al*, 2010; Lito *et al*, 2012, 2014; Poulikakos *et al*, 2010; Solit *et al*, 2006; Yao *et al*, 2015), we distilled decades of structural, biochemical and cell biological work into a single model that provides a self-consistent, unifying picture of RAFi and MEKi resistance in BRAF mutant cancers. The model also correctly captures detailed biochemical properties of MAPK inhibitors without having been explicitly trained on the respective biochemistry: one example is the profound inhibition of EGF activated pERK signaling by type II RAF inhibitors. These features of our model increase confidence that it is a useful and realtively faithful representation of the essential features of intracellular biochemistry. However, some subtleties of MAPK regulation are missing from the model, including the kinase-kinase interactions mediated by KSR scaffolding proteins. The relevance of scaffolding becomes evident in BRAF^V600E^ NRAS^Q61K^ cells exposed to multiple kinase inhibitors. It will be straightforward to add to these features to the model as additional training data becomes available.

In the treatment of melanoma, RAF and MEK inhibitors are used in combination, which is consistent with the more general use of drug combinations to improve reduce resistance to targeted therapy (Lehár *et al*, 2009). Simulation represents an effective way to investigate mechanisms of drug interaction (Fröhlich *et al*, 2018; Yuan *et al*, 2020) and it has been postulated, on theoretical grounds, that inhibition of enzymes acting sequentially in a pathway is a means to achieve synergistic drug interaction (Yin *et al*, 2014; Fitzgerald *et al*, 2006). However, both data and modeling show that the activities of RAF and MEK inhibitors in BRAF^V600E^ cells are additive over the great majority of the dose-response landscape. In those rare conditions in which drug synergy or antagonism is observed, analysis suggests that transcriptional feedback and allosteric interaction – rather than the presence of a serial network motif *per se* – is responsible for drug interaction.

MARM2.0 demonstrates how adaptive drug resistance in BRAF^V600E^ melanoma cells arises from the co- existence in cells of two functionally distinct MAPK reaction channels. Signaling in one channel is initiated by the constitutive activity of oncogenic BRAF^V600E^ and signaling in the other by RAS, which is in turn activated by RTKs. While it is conceptually convenient to depict the BRAF^V600E^ and RAS channels as two different “pathways” (something we do for convenience in **Figure 4**) the actual mechanisms in cells involve shared molecular components: the two reaction channels comprise transient oligomers that involve similar, if not identical, proteins whose dynamic assembly and disassembly allows component exchange. Depending on conditions, one or the other reaction channels can be dominant in regulating ERK, but the two channels can also operate concurrently, masking each other’s activity. For example, in EGF-treated cells, pMEK levels remain roughly constant over a 10^5^-fold range of RAFi because signaling transitions from the BRAF^V600E^ to the RAS channel. The BRAF^V600E^ and RAS channels also influence each other directly, via retroactivity, and indirectly via control over the synthesis of feedback regulators. An additional feature of these reactions is that they operate on multiple time scales; in the case of the RAS channel this includes: (i) a time scale of seconds to minutes involving post-translational modifications and the direct action of inhibitory drugs (ii) a time scale of tens of minutes involving receptor internalization, degradation and recycling and (iii) a time scale of hours involving changes in the levels of negative feedback regulators such as DUSPS and SPRY. Time-scale separation between signal propagation and transcriptional rewiring is necessary for pulsatile signaling to escape from negative feedback and homeostatic control.

Methodological innovation in the current paper focuses on combining rule-based modeling based on PySB and BNG with thermodynamic formalisms that exploit the fact that protein-protein and protein-small molecule binding and unbinding events do not consume energy. This builds on the work of Kholodenko on energy-balanced ODE models (Kholodenko, 2015) while creating a general-purpose framework for programmatically generating model families that make model calibration more efficient. Submodels were generated in PySB to optimally exploit the perturbational structure of the training data (the inclusion or not of drugs and growth factors in each experiment) and combined this with multi-model parameter estimation in the pyPESTO toolbox to substantially accelerate model training, an important consideration with large ODE models and complex training data. Furthermore, PySB/BNG enabled us to implement a labelling scheme for causal network decomposition that traces how species such as activated ERK (e.g. pERK) are generated by converging upstream reaction channels. Analogous generation and analysis of causal traces (“stories”) has been described in agent-based modeling (Boutillier *et al*, 2018) and their adaptation to the MARM2.0 ODE model was essential for formalizing the concept of network rewiring. These and other methods are generally applicable to other models in PySB (Lopez *et al*, 2013) although, in its current implementation, labeling is only designed to trace a sequence of activating events.

Using energies (ΔG and ΔΔG values), rather than kinetic rates, to describe molecular interactions is a more natural and extendable framework for parameterizing biochemical models. Energies can be estimated from structural studies, from mass-spectrometry measurements (Mason & Covert, 2018; de Souza & Picotti, 2020), and increasingly from folding and docking algorithms that combine biophysical understanding of protein structure with deep learning (AlQuraishi & Sorger, 2021; Jumper *et al*, 2021). Approximate energy values can also mitigate the parametric uncertainty that is a pervasive to dynamical models: We anticipate that use of measured or estimated energy values will, in the future, make it possible to place fairly tight priors on parameter values during model calibration, generating more predictive and more interpretable models. Moreover, the use of energy methods promises to bridge the gap between fine-grained atomistic and structural data on single proteins and protein complexes and the more coarse-grained description of biomolecular interactions that are used for dynamical modelling of cellular networks. We anticipate that this will facilitate the multi-scale analysis of allosteric interactions in the assembly of multi- protein(-drug) complexes, and the identification of non-obvious emergent properties.

## MATERIAL AND METHODS

All code that was used to calibrate the model, make predictions and generate figures is available at https://github.com/labsyspharm/marm2-supplement

### Cell lines and tissue culture

The following cell lines were used in this study with their source indicated in parenthesis: A375 (ATCC), A375 with CRISPRa EGFR overexpression (constructed from ATCC stock as reported in (Gerosa *et al*, 2020)), HT29 (Merrimack Pharmaceuticals) and A375 with doxycycline-inducible NRAS^Q61K^ (Yao *et al*, 2015) (provided by Neal Rosen’s lab at Memorial Sloan Kettering Cancer Center). A375 cells were grown in Dulbecco’s modified eagle medium with 4.5 g/l D-glucose, 4 mM L-glutamine, and 1 mM sodium pyruvate (DMEM) (Corning), supplemented with 5% FBS. HT29 cells were grown in RPMI media with L-glutamine supplemented with 10% FBS (50 mL). All media were supplemented with 1% penicillin and streptomycin. Cells were tested for mycoplasma contamination using the MycoAlert mycoplasma detection kit (Lonza).

### Drugs and growth factors

The following chemicals from MedChem Express were dissolved in dimethyl sulfoxide (DMSO) at 10 mM: vemurafenib, LY3009120, AZ-628, cobimetinib. EGF ligand was obtained from Peprotech (cat# 100-15) and prepared in media supplemented with 0.1% bovine serum albumin.

### Experimental design for combined genetic, ligand and drug perturbations

A375 cells with CRISPRa EGFR overexpression and HT29 cells were treated with the indicated drugs for 24 hrs before being stimulated with EGF or mock-media for 8 hours. A375 cells with doxycycline- inducible NRAS^Q61K^ were treated with doxycycline (10 µM) or mock-media for 24 hours before being treated with the indicated drugs for 24 hours.

### Immunofluorescence staining, quantitation, and analysis for cell cultures

The following primary and conjugated antibodies with specified vendor, animal sources and catalogue numbers were used in immunofluorescence analysis of cells and tissues at the specified dilution ratios: p- ERKT202/Y204 rabbit mAb (Cell Signaling Technology, clone D13.14.4E, Cat# 4370), 1:800; p- MEKS217/221 rabbit mAb (Cell Signaling Technology, Cat# 9121) 1:200, ANTI-FLAG® mouse mAb (Sigma Aldrich, Cat# F1804), 1:1000. Immunofluorescence assays for cultured cells were performed using cells seeded in either 96-well plates (Corning Cat#3603) or 384-well plates (CellCarrier Cat#6007558) for 24 hr and then treated with compounds or ligands either using a Hewlett-Packard D300 Digital Dispenser or by manual dispensing.

Cells were fixed in 4% PFA for 30 min at room temperature (RT) and washed with PBS with 0.1% Tween- 20 (Sigma) (PBS-T), permeabilized in methanol for 10 min at RT, rewashed with PBS-T, and blocked in Odyssey blocking buffer (OBB LI-COR Cat. No. 927401) for 1 hr at RT. Cells were incubated overnight at 4 °C with primary antibodies in OBB. Cells were then stained with rabbit and/or with mouse secondary antibodies from Molecular Probes (Invitrogen) labeled with Alexa Fluor 647 (Cat# A31573) or Alexa Fluor 488 (Cat# A21202) both at 1:2000 dilution. Cells were washed with PBS-T and then PBS and were next incubated in 250 ng/mL Hoechst 33342 and 1:2000 HCS CellMask™ Blue Stain solution (Thermo Scientific) for 20 min. Cells were washed twice with PBS and imaged with a 10× objective using a PerkinElmer Operetta High Content Imaging System. 9-11 sites were imaged in each well for 96-well plates and 4-6 sites for 384-well plates.

Image segmentation, analysis, and signal intensity quantitation were performed using the Columbus software (PerkinElmer). Cytosol and nuclear areas were identified by using two different thresholds on the CellMask™ Blue Stain (low intensity) and Hoechst channels (∼100-fold more intense) were used to define cytosolic and nuclear cell masks, respectively. Cells were identified and enumerated according to successful nuclear segmentation. Unless otherwise specified, immunofluorescence quantifications are average signals of the cytosolic area. In the case of the doxycycline-inducible NRAS^Q61K^ A375 cells, low FLAG intensity was used to remove from analysis cells not expressing FLAG-tagged NRAS^Q61K^: in conditions with doxycycline addition FLAG intensity distributions were markedly bimodal with less than 40% of cells being FLAG negative. Population averages were obtained by averaging values from single- cell segmentation using custom MATLAB 2017a code.

### MultiModel Fitting

To the best of our knowledge, all state-of-the-art toolboxes only allow for fitting of individual models. To allow for simultaneous training of multiple models, we implemented the *AggregatedObjective* class in pyPESTO (https://github.com/ICB-DCM/pyPESTO), which implements the mapping between global optimization variables as well as respective gradients and local model parameter values and gradients.

To generate the individual model variants, we implemented the function *MARM.model.get_model_instance*, which uses PySB to programmatically remove subsets of initial values of EGF, RAFi and MEKi species. For network generation we use BNG to construct differential equations only for species with non-zero concentrations. To further reduce computational burden we implemented the function *MARM.model.cleanup_unused*, which programmatically inspects the generated model and removes unused rules, expressions, parameters and energy patterns.

### Model Calibration

Model optimization was performed using pyPESTO 0.2.10 (https://doi.org/10.5281/zenodo.5827905) with fides (Fröhlich & Sorger) version 0.7.5 (https://doi.org/10.5281/zenodo.6038127) as optimizer and AMICI (Fröhlich *et al*, 2021) version 0.11.25 (https://doi.org/10.5281/zenodo.6025361) as simulation engine. 10^3^ optimization runs were performed using randomly sampled initial parameter values. Parameter boundaries that were used for initial value sampling and as constraints for optimization are provided in the function *MARM.estimation.get_problem* in the supplementary material. Initial parameter values where objective function values could not be evaluated were resampled until evaluation was possible. Optimization convergence settings were 10^-12^ as step-size tolerance and 10^-4^ as absolute gradient tolerance. Objective function gradients were computed using forward sensitivity analysis. Integration was limited to 10^6^ steps and integration tolerances were set to 10^-11^ (absolute) and 10^-9^ (relative). Steady-state tolerances were set to 10^-9^ (absolute) and 10^-7^ (relative).

### Causal Signal Decomposition

To track the causal origin of MEK and ERK phosphorylation, we introduced the concept of reaction channels, which combines ideas from causal pathway analysis (Babur *et al*, 2018) and causal lineage tracing (Boutillier *et al*, 2018): Causal pathway analysis explains the response to a perturbation by identifying a sequence of regulatory mechanisms consistent with experimental data. This is equivalent to finding a path in the causal analysis graph, constructed from the knowledge graph, that connects the perturbation with the experimentally observed quantity (Babur *et al*, 2018; Sharp *et al*, 2019). For rule-based models, the causal analysis graph is equivalent to the influence map. Agent based simulations of rules-based models can be represented as random walks on the influence map (Cristescu *et al*, 2019). Accordingly, causal relationships can be extracted by analyzing the traces of individual agents on the knowledge graph (Boutillier *et al*, 2018). As ODE representations of rule-based models describe the average of a population of agents, individual traces are not available and cannot be used to extract causal properties.

To assign phosphorylated MEK and ERK to the BRAF^V600E^ and RAS channels, we added a ‘channel’ site to MEK and ERK molecules, which acts as a tag to track the source of phosphorylation. Upon phosphorylation of MEK, this channel site is set according to the source of phosphorylation ‘phys’ for phosphorylation by RAS bound RAF dimers and ‘onco’ for phosphorylation by mutated BRAF. The rule-based model formulation ensures that the channel information is propagated on all subsequent modeling steps. For the phosphorylation of ERK, we implement two separate rule variants that set the channel site according to the value channel of the phosphorylating MEK molecule. For both pMEK and pERK, the label is set to ‘NA’ during both dephosphorylation and initialization.

### Signaling Gain

In systems biology, strength of signal transmission is typically quantified as response coefficient or logarithmic gain

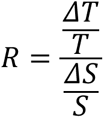

between an input S and an output T at steady-state. However, this definition is not applicable for transient, temporally resolved signals as the response coefficient does not account for the time dimension. As there typically are delays in signal transduction, a pointwise evaluation at individual timepoints does not yield meaningful results.

In signal processing, the gain of linear time invariant systems can be computed as norm of the transfer function G

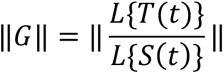

which permits the computation of a gain even for time-resolved inputs *S*(*s*) and outputs *T*(*t*). However, for nonlinear systems, such as the model we developed, a transfer function generally does not exist.

However, we here extend the idea of using functionals such as the Laplace function to map time-resolved input and outputs to scalar values which can then be used to compute the gain. Specifically, we propose the supremum norm

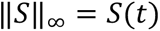

as well as an L1 norm with exponential time transformation

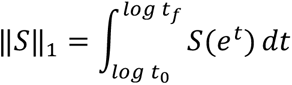

The supremum norm effectively computes the gain evaluated at the peak of the signal, while the L1 norm computes the gain between the area under the curve, where the exponential time transformation aims to avoid problems when signals live on multiple timescales.

The natural scale of gains is the ratio of molecules or concentrations. However, pronounced parameter variability in the estimates for scaling factors, suggested that absolute molecular concentrations were not subject to large uncertainties, which would propagate to these norm estimates. Accordingly, we normalized all gains such that baseline signal transmission had a gain of 1.

To numerically compute supremum and L1 norm, we used 50 log-uniformly spaced time points between 10^-4^ and 10^1^ h. The integral was approximated using the *sklearn.auc* function, which uses the trapezoidal rule.

Despite substantial variability in parameter estimates (**Figure S1**), we found that the variability in qualitative dependence of gain on RAFi and MEKi concentrations is low. We observed the highest variability in the gain from RAS-GTP to physiological pMEK. This is not surprising, as there is no experimental data on RAS-GTP levels. However, the variability appears to primarily affect the absolute levels of signaling gain and less the shape of the dose response curve. Overall, this indicates that our conclusions were not subject to parameter non-identifiability. Moreover, we found that the signaling gain analysis is consistent across different RAFis and MEKis for L1 and L_∞_ norms (**Figure S4**), further corroborating the validity of the approach.

### Predictions for NRAS mutant cell lines

In lack of quantitative measurements of mutant NRAS protein abundances in cell lines with acquired or mutated NRAS, we inferred respective levels from baseline data. In the model, the NRAS mutation was implemented through a constitutive GTP loading reaction that activates RAF independent of upstream receptor activity. Only the rate of this reaction was estimated when retraining on baseline data from respective cell-lines, while all other parameters were kept fixed. For the cell line with acquired NRAS mutation, pERK scaling and offset parameters were simultaneously re-estimated from baseline and naive cell data due to account for difference in data normalization.

### Computation of EC_50_ and EC_max_ values

EC_50_ and EC_max_ were computed by fitting a three-parameter hill function

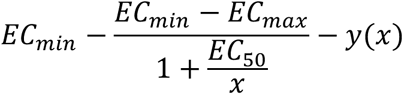

to either experimental data or model simulations, where x are drug concentrations and y are pMEK or pERK levels. *EC_min_* (search interval [0, 2.5], initial 0.5) and *EC_max_* (search interval [0,1.5], initial min (max(*y*(*x_min_*), 0), 2.5) were estimated on a linear scale while *EC*_50_ (search interval [*x_min_*, *x_max_*], initial *x_median_*) was estimated on a logarithmic scale. *scipy.optimize.least_squares* was used for curve fitting.

## ACKNOWLEDGEMENTS

We thank Eduardo Sontag and Pencho Yordanov for helpful discussions. We thank Chris Chidley, Gabriela Sanchez and Sreeram Vallabhaneni for help with experiments. We thank Zhan Yao and Neal Rosen at MSKCC for providing the NRAS^Q61K^ inducible A375 cell line. We thank the Nikon Imaging Center at HMS for assistance with microscopy and the O2 High Performance Compute Cluster for computing support. The work was funded primarily by NCI grant U54-CA225088 (P.K.S.), a Novartis Foundation fellowship to L.G., HFSP grant LT000259/2019-L1 to F.F..

## AUTHOR CONTRIBUTIONS

FF conceived and performed model analysis. LG conceived and performed experiments. LG and FF constructed the model. JM implemented energy support in PySB. PKS supervised the work. FF, LG and PKS wrote the manuscript. All authors reviewed and approved the final version.

---

**Figure S1:**
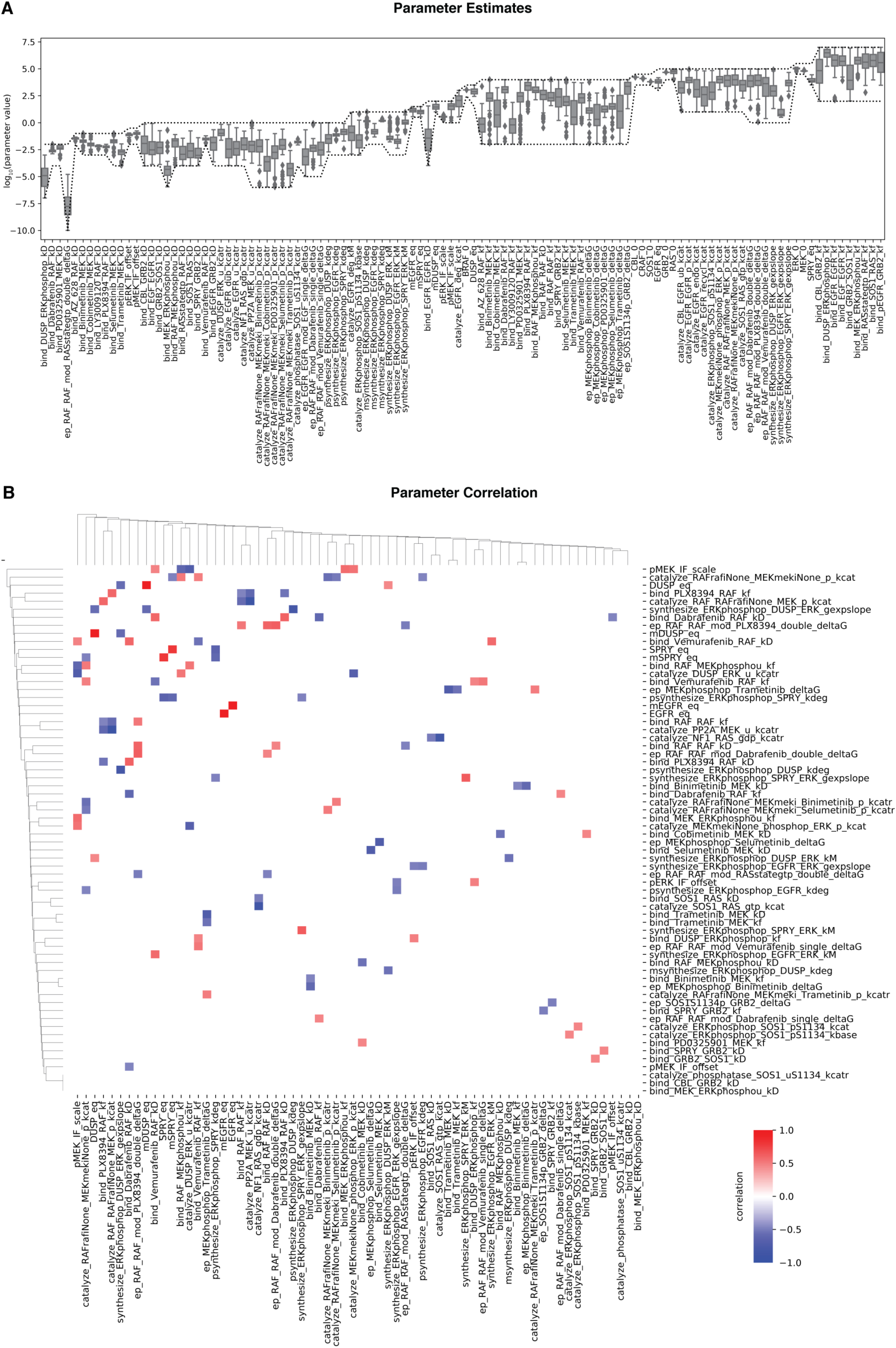
Variability in parameter estimates. **(A)** Boxplot of parameter estimates for best 50 parameter sets. Optimization boundary is indicated as dashed lines. Type of parameters are indicated by suffix: _kD (binding affinity), _offset (background intensity), _kcatr (normalized kcat), _deltaG (thermodynamic parameter), kdeg (degradation rate), kbase (baseline phosphorylation rate), kM (pERK concentration at which 50% activation is achieved), scale (observable scaling), _0 (expression level), _eq (baseline expression level), _kf (binding rate), _kcat (catalytic rate), _gexpslope (RNA synthesis scaling factor). **(B)** Correlation plots of parameter estimates. Only statistically significant (p>0.05) correlations are shown. Coloring shows positive/negative correlation.

**Figure S2:**
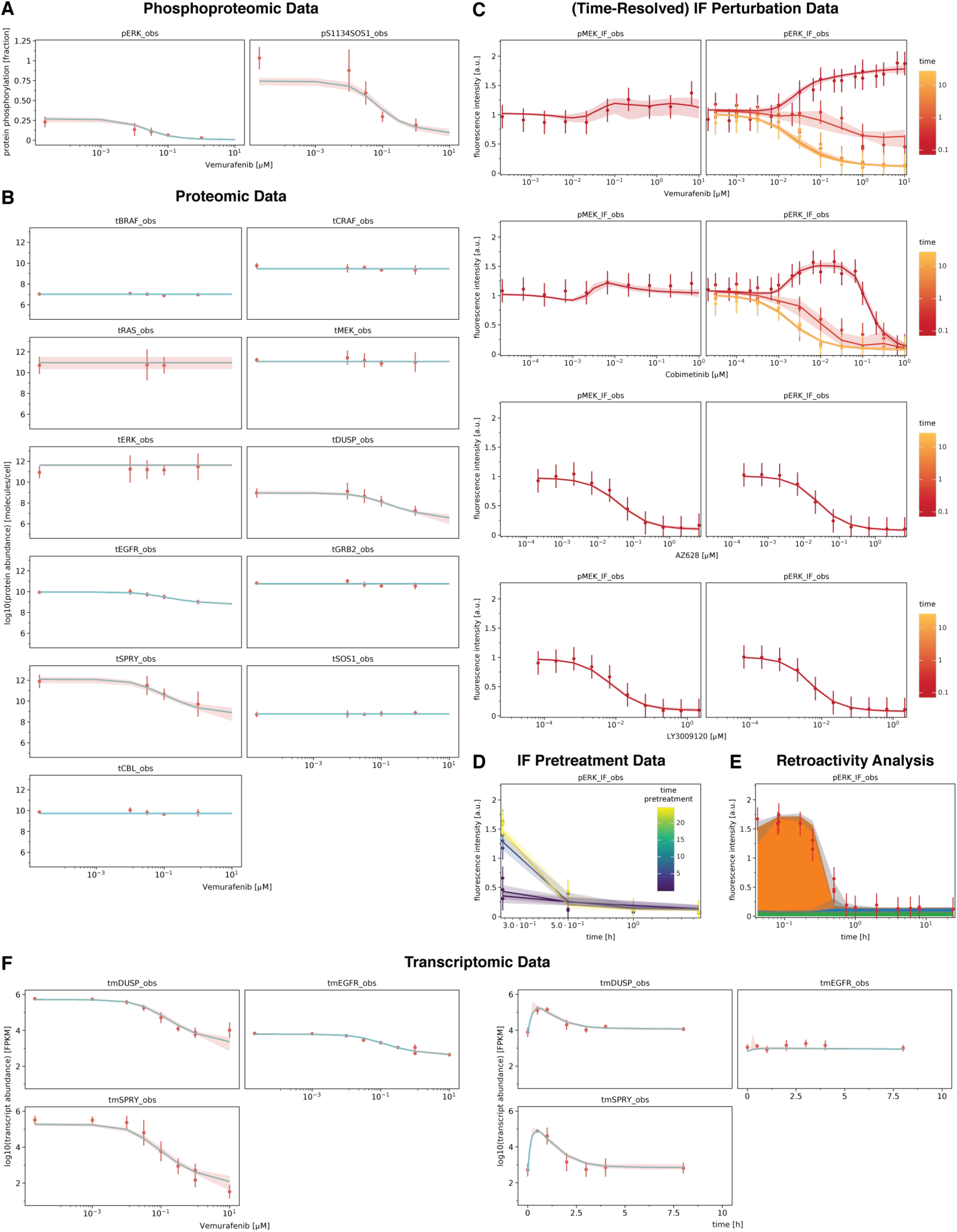
Overview calibrated model simulation and experimental data. Data is shown as point- ranges. Median (over parameter sets) simulations are shown as thick lines. Shading indicates 80% percentiles over parameter sets. **(A)** Phospoproteomic training data (RAFi dose response) **(B)** Proteomic training data (RAFi dose response). **(C)** Additional immunofluorescence data (time resolved RAFi and MEKi dose-response) **(D)** Pretreatment data (timecourse). Pretreatment time indicates the time between drug treatment (1μM vemurafenib) and EGF addition (100ng/ml). **(E)** Causal decomposition of pERK timecourse (1μM vemurafenib) for a modified model in which DUSP can simultaneously bind pERK in the RAS and BRAF^V600E^ channel, preventing retroactivity between channels through DUSP sequestration. **(F)** Transcriptomic training data (RAFi dose response and timecourse)

**Figure S3:**
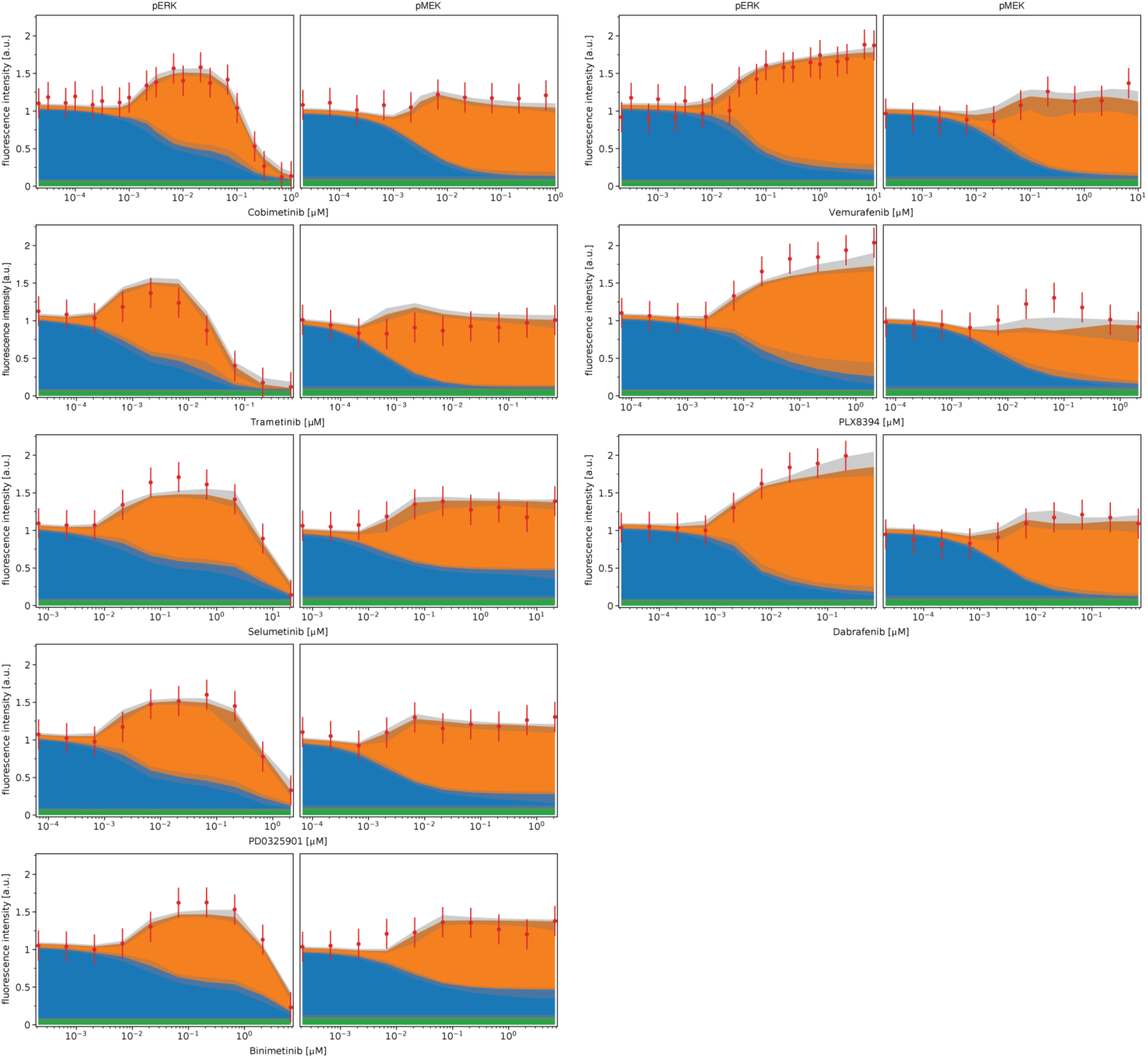
Causal Decomposition of RAS and BRAF^V600E^ channels (extended). Comparison of experimental data and decomposed model simulations at 5 minutes after EGF stimulation for 5 different MEK inhibitors and 3 different RAF inhibitors. Data is shown as point-ranges. Median (over parameter sets) simulations are shown as stacked areas with color corresponding to channels (blue: BRAF^V600E^, orange: RAS). Shading indicates 80% percentiles over parameter sets.

**Figure S4:**
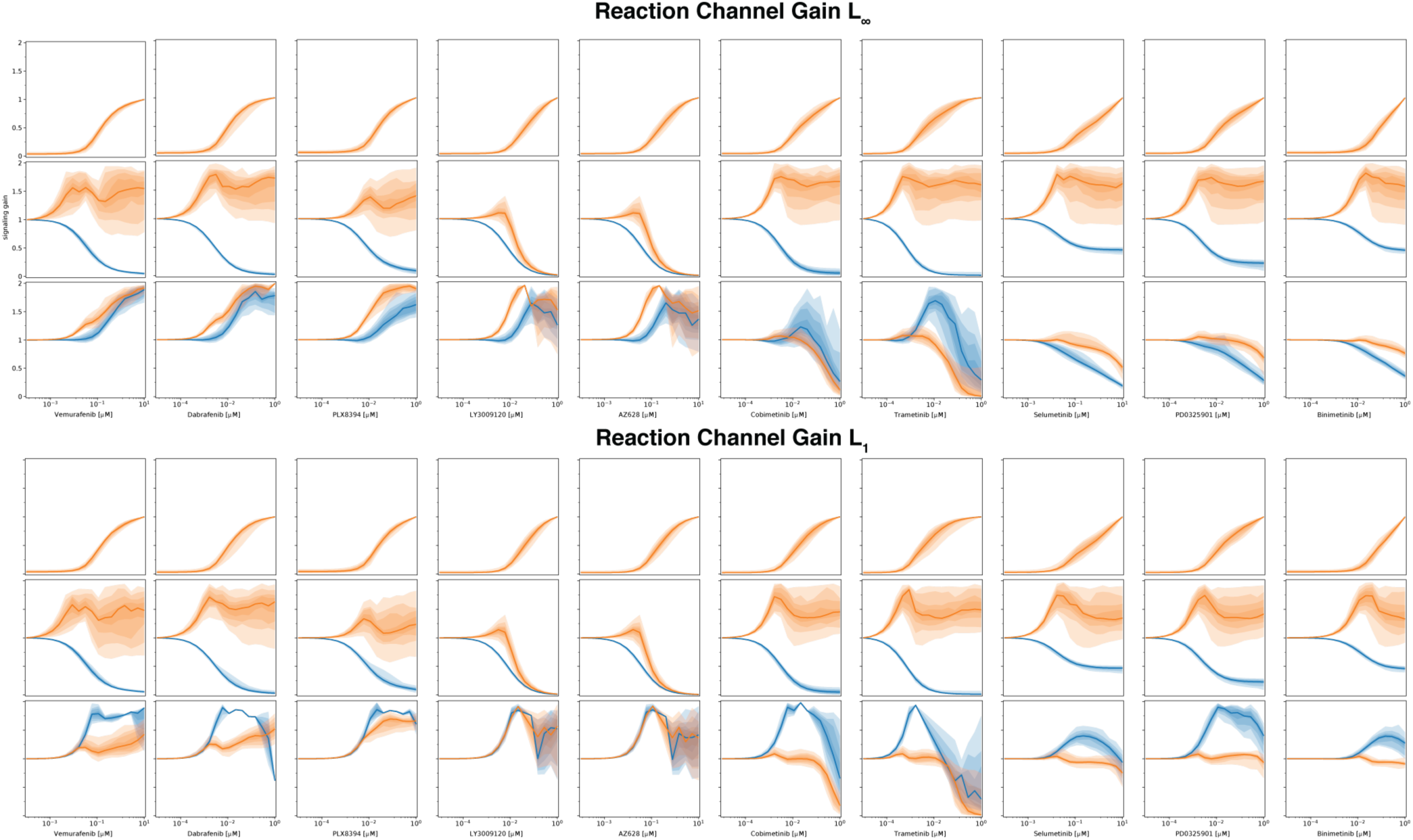
Quantification of signal transduction in RAS and BRAF^V600E^ channels (extended). Quantification of signal transmissions in terms of signaling gain (L_1_ and L∞) along the edges of the simplified network in Figure 4A for different concentrations of 5 different RAF inhibitors and 5 different MEK inhibitors. Color indicates the reaction channel (blue: BRAF^V600E^, orange: RAS). Shading indicates 20, 40, 60 and 80% percentiles over parameter sets.

**Figure S5:**
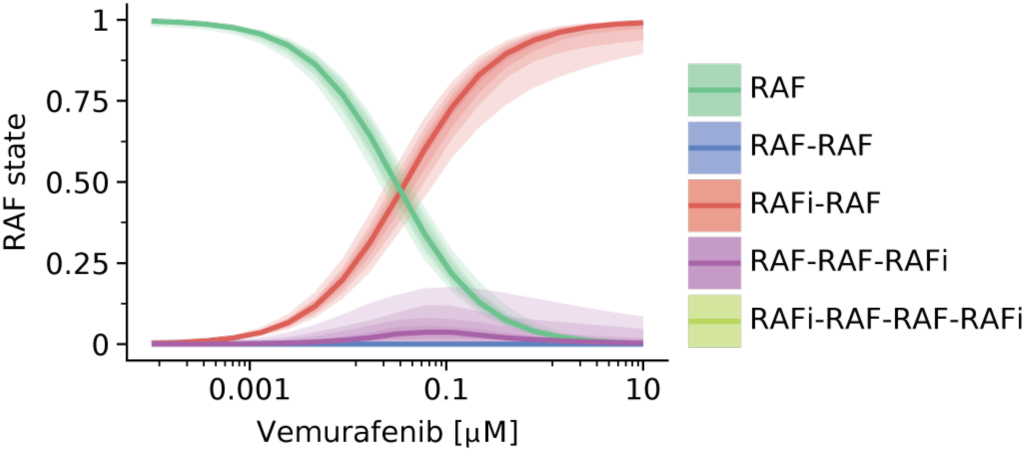
Simulated Assembly RAF-RAFi complexes in response to vemurafenib. Each color corresponds to a different complex. Complex assembly was quantified for RAFi-adapted cells at 5 minutes for dafter EGF stimulation. Shading indicates 20, 40, 60 and 80% percentiles over parameter sets.

**Figure S6:**
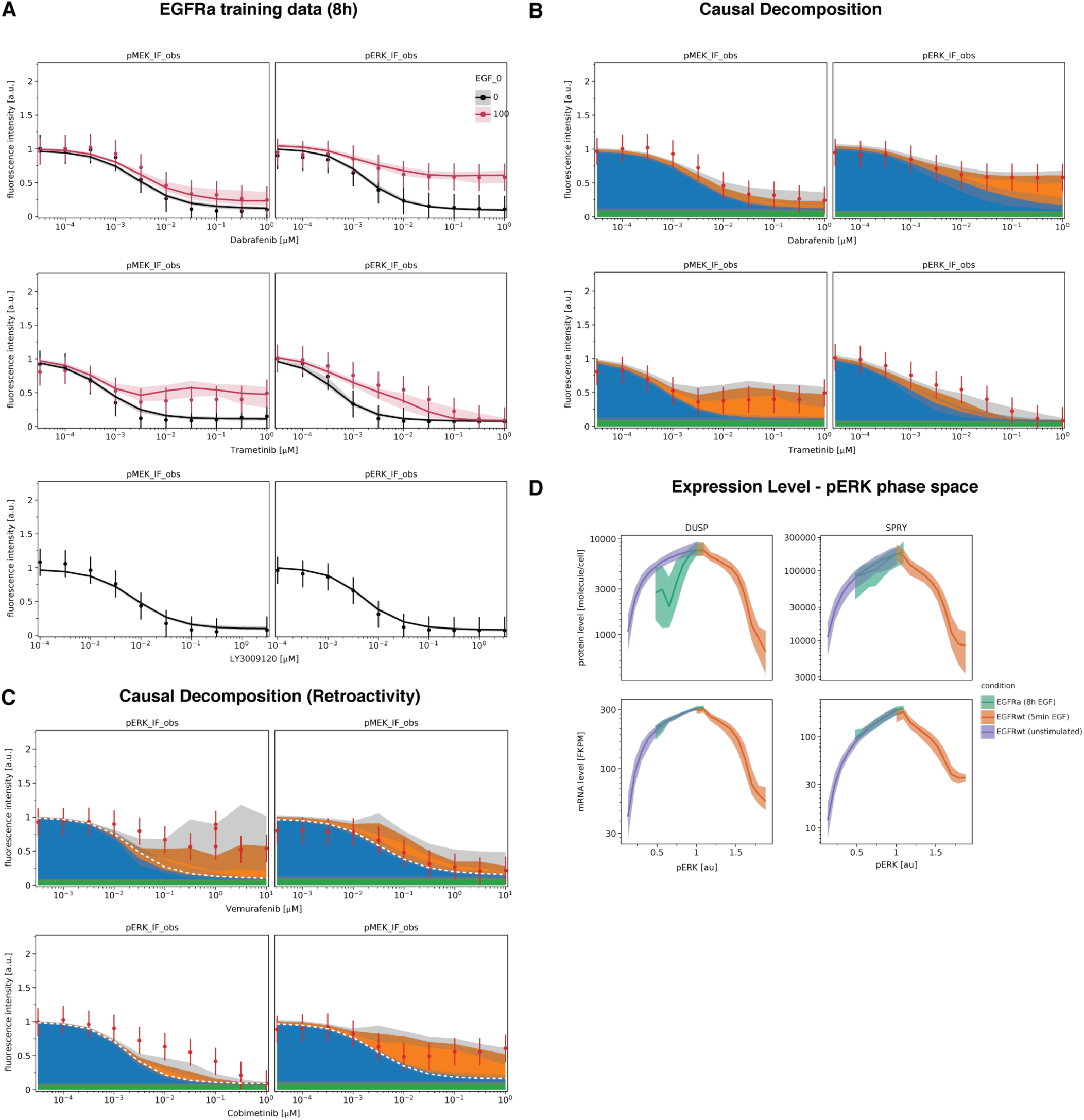
Additional training data for EGFR upregulation and Causal Decomposition. **(A)** Model simulations and experimental data for EGF stimulated and unstimulated conditions. Data is shown as point-ranges. Median (over parameter sets) simulations are shown as thick lines. Shading indicates 80% percentiles over parameter sets. **(B, C)** Comparison of experimental data and decomposed model simulations at 5 minutes after EGF stimulation. Data is shown as point-ranges. Median (over parameter sets) simulations are shown as stacked areas with color corresponding to channels (blue: BRAF^V600E^, orange: RAS). Shading indicates 80% percentiles over parameter sets. C shows causal decomposition of EGF stimulated pMEK and pERK dose response for a modified model in which DUSP can simultaneously bind pERK in the RAS and BRAF^V600E^ channel, preventing retroactivity between channels through DUSP sequestration. Unstimulated baseline indicated by white dashed lines.

**Figure S5:**
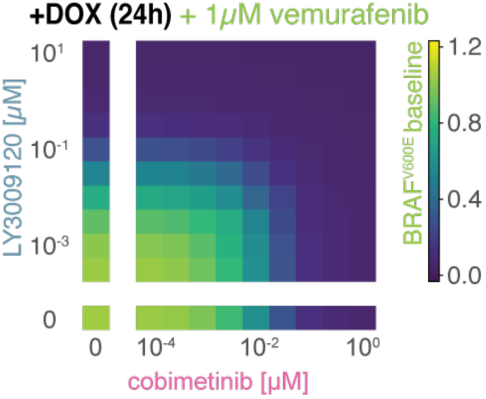
Predicted dose response for combinations of LY3009120 and cobimetinib at 1uM vemurafenib. Simulations were performed for BRAF^V600E^ NRAS^Q61K^ double mutant cells that were adapted to all three drugs.

## REFERENCES

Aldridge BB, Burke JM, Lauffenburger DA & Sorger PK (2006) Physicochemical modelling of cell signalling pathways. Nat Cell Biol 8: 1195–1203

AlQuraishi M & Sorger PK (2021) Differentiable biology: using deep learning for biophysics-based and data-driven modeling of molecular mechanisms. Nat Methods 18: 1169–1180

Alwan HAJ, Zoelen EJJ van & Leeuwen JEM van (2003) Ligand-induced Lysosomal Epidermal Growth Factor Receptor (EGFR) Degradation Is Preceded by Proteasome-dependent EGFR De-ubiquitination *. J Biol Chem 278: 35781–35790

Arrhenius S (1889) Über die Reaktionsgeschwindigkeit bei der Inversion von Rohrzucker durch Säuren. Z Für Phys Chem 4U: 226–248

Babur Ö, Luna A, Korkut A, Durupinar F, Siper MC, Dogrusoz U, Aslan JE, Sander C & Demir E (2018) Causal interactions from proteomic profiles: molecular data meets pathway knowledge. bioRxiv: 258855

Barretina J, Caponigro G, Stransky N, Venkatesan K, Margolin AA, Kim S, Wilson CJ, Lehár J, Kryukov GV, Sonkin D, et al (2012) The Cancer Cell Line Encyclopedia enables predictive modelling of anticancer drug sensitivity. Nature 483: 603–607

Batzer AG, Rotin D, Ureña JM, Skolnik EY & Schlessinger J (1994) Hierarchy of binding sites for Grb2 and Shc on the epidermal growth factor receptor. Mol Cell Biol 14: 5192–5201

Becker V, Schilling M, Bachmann J, Baumann U, Raue A, Maiwald T, Timmer J & Klingmüller U (2010) Covering a broad dynamic range: information processing at the erythropoietin receptor. Science 328: 1404–1408

Blinov ML, Faeder JR, Goldstein B & Hlavacek WS (2004) BioNetGen: software for rule-based modeling of signal transduction based on the interactions of molecular domains. Bioinformatics 20: 3289–3291

Blinov ML, Faeder JR, Goldstein B & Hlavacek WS (2006) A network model of early events in epidermal growth factor receptor signaling that accounts for combinatorial complexity. Biosystems 83: 136–151

Bliss CI (1939) The Toxicity of Poisons Applied Jointly. Ann Appl Biol 26: 585–615

Boutillier P, Maasha M, Li X, Medina-Abarca HF, Krivine J, Feret J, Cristescu I, Forbes AG & Fontana W (2018) The Kappa platform for rule-based modeling. Bioinformatics 34: i583–i592

Burd CE, Liu W, Huynh MV, Waqas MA, Gillahan JE, Clark KS, Fu K, Martin BL, Jeck WR, Souroullas GP, et al (2014) Mutation-Specific RAS Oncogenicity Explains NRAS Codon 61 Selection in Melanoma. Cancer Discov 4: 1418–1429

Burotto M, Chiou VL, Lee J-M & Kohn EC (2014) The MAPK pathway across different malignancies: A new perspective. Cancer 120: 3446–3456

Chen WW, Schoeberl B, Jasper PJ, Niepel M, Nielsen UB, Lauffenburger DA & Sorger PK (2009) Input– output behavior of ErbB signaling pathways as revealed by a mass action model trained against dynamic data. Mol Syst Biol 5

Chis O-T, Banga JR & Balsa-Canto E (2011) Structural identifiability of systems biology models: A critical comparison of methods. PLoS ONE 6: e27755

Chou T-C, Tan Q-H & Sirotnak FM (1993) Quantitation of the synergistic interaction of edatrexate and cisplatin in vitro. Cancer Chemother Pharmacol 31: 259–264

Clarke CN & Kopetz ES (2015) BRAF mutant colorectal cancer as a distinct subset of colorectal cancer: clinical characteristics, clinical behavior, and response to targeted therapies. J Gastrointest Oncol 6: 660–667

Corbalan-Garcia S, Yang SS, Degenhardt KR & Bar-Sagi D (1996) Identification of the mitogen-activated protein kinase phosphorylation sites on human Sos1 that regulate interaction with Grb2. Mol Cell Biol

Cristescu I, Fontana W & Krivine J (2019) Interactions between Causal Structures in Graph Rewriting Systems. Electron Proc Theor Comput Sci 286: 65–78

Davies H, Bignell GR, Cox C, Stephens P, Edkins S, Clegg S, Teague J, Woffendin H, Garnett MJ, Bottomley W, et al (2002) Mutations of the BRAF gene in human cancer. Nature 417: 949

Del Vecchio D, Ninfa AJ & Sontag ED (2008) Modular cell biology: retroactivity and insulation. Mol Syst Biol 4: 161

Dessauges C, Mikelson J, Dobrzyński M, Jacques M-A, Frismantiene A, Gagliardi PA, Khammash M & Pertz O (2021) Optogenetic actuator/biosensor circuits for large-scale interrogation of ERK dynamics identify sources of MAPK signaling robustness. bioRxiv: 2021.07.27.453955 doi:10.1101/2021.07.27.453955 [PREPRINT]

Dhawan NS, Scopton AP & Dar AC (2016) Small molecule stabilization of the KSR inactive state antagonizes oncogenic Ras signalling. Nature 537: 112–116

Ding K-F, Finlay D, Yin H, Hendricks WPD, Sereduk C, Kiefer J, Sekulic A, LoRusso PM, Vuori K, Trent JM, et al (2018) Network Rewiring in Cancer: Applications to Melanoma Cell Lines and the Cancer Genome Atlas Patients. Front Genet 9: 228

English JM & Cobb MH (2002) Pharmacological inhibitors of MAPK pathways. Trends Pharmacol Sci 23: 40–45

Evans MG & Polanyi M (1935) Some applications of the transition state method to the calculation of reaction velocities, especially in solution. Trans Faraday Soc 31: 875–894

Eydgahi H, Chen WW, Muhlich JL, Vitkup D, Tsitsiklis JN & Sorger PK (2013) Properties of cell death models calibrated and compared using Bayesian approaches. Mol Syst Biol 9: 644

Eyring H (1935) The Activated Complex in Chemical Reactions. J Chem Phys 3: 107–115

Faeder JR, Blinov ML, Goldstein B & Hlavacek WS (2005) Combinatorial complexity and dynamical restriction of network flows in signal transduction. Syst Biol 2: 5–15

Fallahi-Sichani M, Becker V, Izar B, Baker GJ, Lin J, Boswell SA, Shah P, Rotem A, Garraway LA & Sorger PK (2017) Adaptive resistance of melanoma cells to RAF inhibition via reversible induction of a slowly dividing de-differentiated state. Mol Syst Biol 13

Fitzgerald JB, Schoeberl B, Nielsen UB & Sorger PK (2006) Systems biology and combination therapy in the quest for clinical efficacy. Nat Chem Biol 2: 458–66

Flaherty KT, Infante JR, Daud A, Gonzalez R, Kefford RF, Sosman J, Hamid O, Schuchter L, Cebon J, Ibrahim N, et al (2012) Combined BRAF and MEK Inhibition in Melanoma with BRAF V600 Mutations. N Engl J Med 367: 1694–1703

Fröhlich F, Kaltenbacher B, Theis FJ & Hasenauer J (2017) Scalable parameter estimation for genome-scale biochemical reaction networks. PLoS Comput Biol 13: 1–18

Fröhlich F, Kessler T, Weindl D, Shadrin A, Schmiester L, Hache H, Muradyan A, Schütte M, Lim J-H, Heinig M, et al (2018) Efficient Parameter Estimation Enables the Prediction of Drug Response Using a Mechanistic Pan-Cancer Pathway Model. Cell Syst 7: 567–579.e6

Fröhlich F, Loos C & Hasenauer J (2019) Scalable Inference of Ordinary Differential Equation Models of Biochemical Processes. In Gene Regulatory Networks: Methods and Protocols, Sanguinetti G & Huynh-Thu VA (eds) pp 385–422. New York, NY: Springer

Fröhlich F & Sorger PK Fides: Reliable Trust-Region Optimization for Parameter Estimation of Ordinary Differential Equation Models. bioRxiv

Fröhlich F, Theis FJ & Hasenauer J (2014) Uncertainty analysis for non-identifiable dynamical systems: Profile likelihoods, bootstrapping and more. In Proceedings of the 12th International Conference on Computational Methods in Systems Biology (CMSB 2014), Manchester, UK, Mendes P Dada JO & Smallbone KO (eds) pp 61–72. Springer International Publishing Switzerland

Fröhlich F, Thomas P, Kazeroonian A, Theis FJ, Grima R & Hasenauer J (2016) Inference for stochastic chemical kinetics using moment equations and system size expansion. PLoS Comput Biol 12: e1005030

Fröhlich F, Weindl D, Schälte Y, Pathirana D, Paszkowski Ł, Lines GT, Stapor P & Hasenauer J (2021) AMICI: High-Performance Sensitivity Analysis for Large Ordinary Differential Equation Models. Bioinformatics 37: 3676–3677

Gao Y, Chang MT, McKay D, Na N, Zhou B, Yaeger R, Torres NM, Muniz K, Drosten M, Barbacid M, et al (2018) Allele-Specific Mechanisms of Activation of MEK1 Mutants Determine Their Properties. Cancer Discov 8: 648–661

Gawthrop PJ & Crampin EJ (2017) Energy-based analysis of biomolecular pathways. Proc R Soc Math Phys Eng Sci 473: 20160825

Gerosa L, Chidley C, Fröhlich F, Sanchez G, Lim SK, Muhlich J, Chen J-Y, Vallabhaneni S, Baker GJ, Schapiro D, et al (2020) Receptor-Driven ERK Pulses Reconfigure MAPK Signaling and Enable Persistence of Drug-Adapted BRAF-Mutant Melanoma Cells. Cell Syst 11: 478–494.e9

Goldbeter A & Koshland DE (1981) An amplified sensitivity arising from covalent modification in biological systems. Proc Natl Acad Sci U S A 78: 6840–6844

Gollub MG, Kaltenbach H-M & Stelling J (2021) Probabilistic thermodynamic analysis of metabolic networks. Bioinformatics 37: 2938–2945

Gutenkunst RN, Waterfall JJ, Casey FP, Brown KS, Myers CR & Sethna JP (2007) Universally sloppy parameter sensitivities in systems biology models. PLoS Comput Biol 3: 1871–1878

Haling JR, Sudhamsu J, Yen I, Sideris S, Sandoval W, Phung W, Bravo BJ, Giannetti AM, Peck A, Masselot A, et al (2014) Structure of the BRAF-MEK Complex Reveals a Kinase Activity Independent Role for BRAF in MAPK Signaling. Cancer Cell 26: 402–413

Hall-Jackson CA, Eyers PA, Cohen P, Goedert M, Tom Boyle F, Hewitt N, Plant H & Hedge P (1999) Paradoxical activation of Raf by a novel Raf inhibitor. Chem Biol 6: 559–568

Harris LA, Hogg JS, Tapia J-J, Sekar JAP, Gupta S, Korsunsky I, Arora A, Barua D, Sheehan RP & Faeder JR (2016) BioNetGen 2.2: advances in rule-based modeling. Bioinforma Oxf Engl 32: 3366–3368

Hatzivassiliou G, Haling JR, Chen H, Song K, Price S, Heald R, Hewitt JFM, Zak M, Peck A, Orr C, et al (2013) Mechanism of MEK inhibition determines efficacy in mutant KRAS- versus BRAF-driven cancers. Nature 501: 232–236

Hatzivassiliou G, Song K, Yen I, Brandhuber BJ, Anderson DJ, Alvarado R, Ludlam MJC, Stokoe D, Gloor SL, Vigers G, et al (2010) RAF inhibitors prime wild-type RAF to activate the MAPK pathway and enhance growth. Nature 464: 431–435

Henry JR, Kaufman MD, Peng S-B, Ahn YM, Caldwell TM, Vogeti L, Telikepalli H, Lu W-P, Hood MM, Rutkoski TJ, et al (2015) Discovery of 1-(3,3-Dimethylbutyl)-3-(2-fluoro-4-methyl-5-(7-methyl-2- (methylamino)pyrido[2,3-d]pyrimidin-6-yl)phenyl)urea (LY3009120) as a Pan-RAF Inhibitor with Minimal Paradoxical Activation and Activity against BRAF or RAS Mutant Tumor Cells. J Med Chem 58: 4165–4179

Hlavacek WS, Faeder JR, Blinov ML, Posner RG, Hucka M & Fontana W (2006) Rules for Modeling Signal-Transduction Systems. Sci STKE 2006

Hogg JS (2013) Advances in Rule-based Modeling: Compartments, Energy, and Hybrid Simulation, with Application to Sepsis and Cell Signaling. undefined

Honorato-Zimmer R, Harmer R & Danos V (2015) Thermodynamic graph-rewriting. Log Methods Comput Sci Volume 11, Issue 2

Hunter T (2000) Signaling--2000 and beyond. Cell 100: 113–127

Hyttinen A, Eberhardt F & Hoyer PO (2012) Learning Linear Cyclic Causal Models with Latent Variables. J Mach Learn Res 13: 3387–3439

Jumper J, Evans R, Pritzel A, Green T, Figurnov M, Ronneberger O, Tunyasuvunakool K, Bates R, Žídek A, Potapenko A, et al (2021) Highly accurate protein structure prediction with AlphaFold. Nature: 1– 11

Kamioka Y, Yasuda S, Fujita Y, Aoki K & Matsuda M (2010) Multiple Decisive Phosphorylation Sites for the Negative Feedback Regulation of SOS1 via ERK *. J Biol Chem 285: 33540–33548

Karoulia Z, Gavathiotis E & Poulikakos PI (2017) New perspectives for targeting RAF kinase in human cancer. Nat Rev Cancer 17: 676–691

Karoulia Z, Wu Y, Ahmed TA, Xin Q, Bollard J, Krepler C, Wu X, Zhang C, Bollag G, Herlyn M, et al (2016) An Integrated Model of RAF Inhibitor Action Predicts Inhibitor Activity against Oncogenic BRAF Signaling. Cancer Cell 30: 485–498

Kebebew E, Weng J, Bauer J, Ranvier G, Clark OH, Duh Q-Y, Shibru D, Bastian B & Griffin A (2007) The Prevalence and Prognostic Value of BRAF Mutation in Thyroid Cancer. Ann Surg 246: 466–471

Kholodenko BN (2015) Drug Resistance Resulting from Kinase Dimerization Is Rationalized by Thermodynamic Factors Describing Allosteric Inhibitor Effects. Cell Rep 12: 1939–1949

Kholodenko BN, Demin OV, Moehren G & Hoek JB (1999) Quantification of Short Term Signaling by the Epidermal Growth Factor Receptor. J Biol Chem 274: 30169–30181

Kiyatkin A, Rosenburgh IK van A van, Klein DE & Lemmon MA (2020) Kinetics of receptor tyrosine kinase activation define ERK signaling dynamics. Sci Signal 13

Kleiman LB, Maiwald T, Conzelmann H, Lauffenburger DA & Sorger PK (2011) Rapid phospho-turnover by receptor tyrosine kinases impacts downstream signaling and drug binding. Mol Cell 43: 723–737

Klosin A, Oltsch F, Harmon T, Honigmann A, Jülicher F, Hyman AA & Zechner C (2020) Phase separation provides a mechanism to reduce noise in cells. Science 367: 464–468

Kreutz C, Raue A & Timmer J (2012) Likelihood based observability analysis and confidence intervals for predictions of dynamic models. BMC Syst Biol 6

Lao D-H, Chandramouli S, Yusoff P, Fong CW, Saw TY, Tai LP, Yu CY, Leong HF & Guy GR (2006) A Src Homology 3-binding Sequence on the C Terminus of Sprouty2 Is Necessary for Inhibition of the Ras/ERK Pathway Downstream of Fibroblast Growth Factor Receptor Stimulation *. J Biol Chem 281: 29993–30000

Lavoie H, Gagnon J & Therrien M (2020) ERK signalling: a master regulator of cell behaviour, life and fate. Nat Rev Mol Cell Biol 21: 607–632

Lavoie H & Therrien M (2015) Regulation of RAF protein kinases in ERK signalling. Nat Rev Mol Cell Biol 16: 281–298

Lee MJ, Ye AS, Gardino AK, Heijink AM, Sorger PK, MacBeath G & Yaffe MB (2012) Sequential application of anticancer drugs enhances cell death by rewiring apoptotic signaling networks. Cell 149: 780–794

Lehár J, Krueger AS, Avery W, Heilbut AM, Johansen LM, Price ER, Rickles RJ, Short Iii GF, Staunton JE, Jin X, et al (2009) Synergistic drug combinations tend to improve therapeutically relevant selectivity. Nat Biotechnol 27: 659–666

Lehár J, Zimmermann GR, Krueger AS, Molnar RA, Ledell JT, Heilbut AM, Short GF, Giusti LC, Nolan GP, Magid OA, et al (2007) Chemical combination effects predict connectivity in biological systems. Mol Syst Biol 3

Lemmon MA & Schlessinger J (2010) Cell signaling by receptor-tyrosine kinases. Cell 141: 1117–1134

Lito P, Pratilas CA, Joseph EW, Tadi M, Halilovic E, Zubrowski M, Huang A, Wong WL, Callahan MK, Merghoub T, et al (2012) Relief of Profound Feedback Inhibition of Mitogenic Signaling by RAF Inhibitors Attenuates their Activity in BRAFV600E Melanomas. Cancer Cell 22: 668–682

Lito P, Rosen N & Solit DB (2013) Tumor adaptation and resistance to RAF inhibitors. Nat Med 19: 1401

Lito P, Saborowski A, Yue J, Solomon M, Joseph E, Gadal S, Saborowski M, Kastenhuber E, Fellmann C, Ohara K, et al (2014) Disruption of CRAF-Mediated MEK Activation Is Required for Effective MEK Inhibition in KRAS Mutant Tumors. Cancer Cell 25: 697–710

Loewe S (1928) Die quantitativen Probleme der Pharmakologie. Ergeb Physiol 27: 47–187

Long GV, Fung C, Menzies AM, Pupo GM, Carlino MS, Hyman J, Shahheydari H, Tembe V, Thompson JF, Saw RP, et al (2014) Increased MAPK reactivation in early resistance to dabrafenib/trametinib combination therapy of BRAF-mutant metastatic melanoma. Nat Commun 5: 1–9

Lopez CF, Muhlich JL, Bachman JA & Sorger PK (2013) Programming biological models in Python using PySB. Mol Syst Biol 9: 646

Lund KA, Opresko LK, Starbuck C, Walsh BJ & Wiley HS (1990) Quantitative analysis of the endocytic system involved in hormone-induced receptor internalization. J Biol Chem 265: 15713–15723

Marin-Bejar O, Rogiers A, Dewaele M, Femel J, Karras P, Pozniak J, Bervoets G, Raemdonck NV, Pedri D, Swings T, et al (2021) Evolutionary predictability of genetic versus nongenetic resistance to anticancer drugs in melanoma. Cancer Cell 39: 1135–1149.e8

Mason JC & Covert MW (2018) An energetic reformulation of kinetic rate laws enables scalable parameter estimation for biochemical networks. J Theor Biol

Mooij JM, Janzing D & Schölkopf B (2013) From Ordinary Differential Equations to Structural Causal Models: the deterministic case. *ArXiv13047920 Cs Stat*

Noeparast A, Giron P, De Brakeleer S, Eggermont C, De Ridder U, Teugels E & De Grève J (2018) Type II RAF inhibitor causes superior ERK pathway suppression compared to type I RAF inhibitor in cells expressing different BRAF mutant types recurrently found in lung cancer. Oncotarget 9: 16110–16123

Nusinow DP, Szpyt J, Ghandi M, Rose CM, McDonald ER, Kalocsay M, Jané-Valbuena J, Gelfand E, Schweppe DK, Jedrychowski M, et al (2020) Quantitative Proteomics of the Cancer Cell Line Encyclopedia. Cell 180: 387–402.e16

Olivier BG, Rohwer JM & Hofmeyr J-HS (2005) Modelling cellular systems with PySCeS. Bioinf 21: 560–561

Ollivier JF, Shahrezaei V & Swain PS (2010) Scalable Rule-Based Modelling of Allosteric Proteins and Biochemical Networks. PLOS Comput Biol 6: e1000975

Oren Y, Tsabar M, Cuoco MS, Amir-Zilberstein L, Cabanos HF, Hütter J-C, Hu B, Thakore PI, Tabaka M, Fulco CP, et al (2021) Cycling cancer persister cells arise from lineages with distinct programs. Nature 596: 576–582

Pearl J & Dechter R (2013) Identifying Independencies in Causal Graphs with Feedback. *ArXiv13023595 Cs*

Peng S-B, Henry JR, Kaufman MD, Lu W-P, Smith BD, Vogeti S, Rutkoski TJ, Wise S, Chun L, Zhang Y, et al (2015) Inhibition of RAF Isoforms and Active Dimers by LY3009120 Leads to Anti-tumor Activities in RAS or BRAF Mutant Cancers. Cancer Cell 28: 384–398

Pino GLG-D, Li K, Park E, Schmoker AM, Ha BH & Eck MJ (2021) Allosteric MEK inhibitors act on BRAF/MEK complexes to block MEK activation. Proc Natl Acad Sci 118

Poulikakos PI, Persaud Y, Janakiraman M, Kong X, Ng C, Moriceau G, Shi H, Atefi M, Titz B, Gabay MT, et al (2011) RAF inhibitor resistance is mediated by dimerization of aberrantly spliced BRAF(V600E). Nature 480: 387–390

Poulikakos PI, Zhang C, Bollag G, Shokat KM & Rosen N (2010) RAF inhibitors transactivate RAF dimers and ERK signalling in cells with wild-type BRAF. Nature 464: 427–430

Pratilas CA, Taylor BS, Ye Q, Viale A, Sander C, Solit DB & Rosen N (2009) (V600E)BRAF is associated with disabled feedback inhibition of RAF-MEK signaling and elevated transcriptional output of the pathway. Proc Natl Acad Sci U S A 106: 4519–4524

Prior IA, Lewis PD & Mattos C (2012) A Comprehensive Survey of Ras Mutations in Cancer. Cancer Res 72: 2457–2467

Raue A, Kreutz C, Maiwald T, Klingmüller U & Timmer J (2011) Addressing parameter identifiability by model-based experimentation. IET Syst Biol 5: 120–130

Reddy RJ, Gajadhar AS, Swenson EJ, Rothenberg DA, Curran TG & White FM (2016) Early signaling dynamics of the epidermal growth factor receptor. Proc Natl Acad Sci 113: 3114–3119

Resat H, Ewald JA, Dixon DA & Wiley HS (2003) An Integrated Model of Epidermal Growth Factor Receptor Trafficking and Signal Transduction. Biophys J 85: 730–743

Roskoski R (2016) Classification of small molecule protein kinase inhibitors based upon the structures of their drug-enzyme complexes. Pharmacol Res 103: 26–48

Rukhlenko OS, Khorsand F, Krstic A, Rozanc J, Alexopoulos LG, Rauch N, Erickson KE, Hlavacek WS, Posner RG, Gómez-Coca S, et al (2018) Dissecting RAF Inhibitor Resistance by Structure-based Modeling Reveals Ways to Overcome Oncogenic RAS Signaling. Cell Syst 7: 161–179.e14

Russo M, Crisafulli G, Sogari A, Reilly NM, Arena S, Lamba S, Bartolini A, Amodio V, Magrì A, Novara L, et al (2019) Adaptive mutability of colorectal cancers in response to targeted therapies. Science 366: 1473–1480

Samatar AA & Poulikakos PI (2014) Targeting RAS–ERK signalling in cancer: promises and challenges. Nat Rev Drug Discov 13: 928–942

Sanchez-Vega F, Mina M, Armenia J, Chatila WK, Luna A, La KC, Dimitriadoy S, Liu DL, Kantheti HS, Saghafinia S, et al (2018) Oncogenic Signaling Pathways in The Cancer Genome Atlas. Cell 173: 321–337.e10

Sauro HM (2008) Modularity defined. Mol Syst Biol 4: 166

Schöberl B, Pace EA, Fitzgerald JB, Harms BD, Xu L, Nie L, Linggi B, Kalra A, Paragas V, Bukhalid R, et al (2009) Therapeutically targeting ErbB3: A key node in ligand-induced activation of the ErbB receptor-PI3K axis. Sci Signal 2: ra31

Schuh L, Saint-Antoine M, Sanford EM, Emert BL, Singh A, Marr C, Raj A & Goyal Y (2020) Gene Networks with Transcriptional Bursting Recapitulate Rare Transient Coordinated High Expression States in Cancer. Cell Syst 10: 363–378.e12

Sekar JAP, Hogg JS & Faeder JR (2016) Energy-based modeling in BioNetGen. In 2016 IEEE International Conference on Bioinformatics and Biomedicine (BIBM) pp 1460–1467.

Shaffer SM, Dunagin MC, Torborg SR, Torre EA, Emert B, Krepler C, Beqiri M, Sproesser K, Brafford PA, Xiao M, et al (2017) Rare cell variability and drug-induced reprogramming as a mode of cancer drug resistance. Nature 546: 431–435

Sharp R, Pyarelal A, Gyori B, Alcock K, Laparra E, Valenzuela-Escárcega MA, Nagesh A, Yadav V, Bachman J, Tang Z, et al (2019) Eidos, INDRA, & Delphi: From Free Text to Executable Causal Models. In Proceedings of the 2019 Conference of the North American Chapter of the Association for Computational Linguistics (Demonstrations) pp 42–47. Minneapolis, Minnesota: Association for Computational Linguistics

Shi H, Hugo W, Kong X, Hong A, Koya RC, Moriceau G, Chodon T, Guo R, Johnson DB, Dahlman KB, et al (2014) Acquired resistance and clonal evolution in melanoma during BRAF inhibitor therapy. Cancer Discov 4: 80–93

Sneddon MW, Faeder JR & Emonet T (2011) Efficient modeling, simulation and coarse-graining of biological complexity with NFsim. Nat Methods 8: 177–183

Solit DB, Garraway LA, Pratilas CA, Sawai A, Getz G, Basso A, Ye Q, Lobo JM, She Y, Osman I, et al (2006) BRAF mutation predicts sensitivity to MEK inhibition. Nature 439: 358–362

de Souza N & Picotti P (2020) Mass spectrometry analysis of the structural proteome. Curr Opin Struct Biol 60: 57–65

Spirtes PL (2013) Directed Cyclic Graphical Representations of Feedback Models. *ArXiv13024982 Cs*

Städter P, Schälte Y, Schmiester L, Hasenauer J & Stapor PL (2021) Benchmarking of numerical integration methods for ODE models of biological systems. Sci Rep 11: 2696

Stapor P, Weindl D, Ballnus B, Hug S, Loos C, Fiedler A, Krause S, Hroß S, Fröhlich F & Hasenauer J (2018) PESTO: Parameter EStimation TOolbox. Bioinforma Oxf Engl 34: 705–707

Starbuck C & Lauffenburger DA (1992) Mathematical model for the effects of epidermal growth factor receptor trafficking dynamics on fibroblast proliferation responses. Biotechnol Prog 8: 132–143

Sullivan RJ & Flaherty K (2012) MAP kinase signaling and inhibition in melanoma. Oncogene 32: 2373

Tsai C-J & Nussinov R (2014) A Unified View of “How Allostery Works”. PLOS Comput Biol 10: e1003394

Tutuka CSA, Andrews MC, Mariadason JM, Ioannidis P, Hudson C, Cebon J & Behren A (2017) PLX8394, a new generation BRAF inhibitor, selectively inhibits BRAF in colonic adenocarcinoma cells and prevents paradoxical MAPK pathway activation. Mol Cancer 16: 112

Ullrich A & Schlessinger J (1990) Signal transduction by receptors with tyrosine kinase activity. Cell 61: 203–212

Villaverde AF, Fröhlich F, Weindl D, Hasenauer J & Banga JR (2019) Benchmarking optimization methods for parameter estimation in large kinetic models. Bioinformatics 35: 830–838

Wagle N, Emery C, Berger MF, Davis MJ, Sawyer A, Pochanard P, Kehoe SM, Johannessen CM, MacConaill LE, Hahn WC, et al (2011) Dissecting Therapeutic Resistance to RAF Inhibition in Melanoma by Tumor Genomic Profiling. J Clin Oncol 29: 3085–3096

Wegscheider R (1911) Über simultane Gleichgewichte und die Beziehungen zwischen Thermodynamik und Reactionskinetik homogener Systeme. Monatshefte Für Chem Verwandte Teile Anderer Wiss 32: 849–906

Wei Q, Qian Y, Yu J & Wong CC (2020) Metabolic rewiring in the promotion of cancer metastasis: mechanisms and therapeutic implications. Oncogene 39: 6139–6156

Wieland F-G, Hauber AL, Rosenblatt M, Tönsing C & Timmer J (2021) On structural and practical identifiability. Curr Opin Syst Biol 25: 60–69

Wiley HS (1988) Anomalous binding of epidermal growth factor to A431 cells is due to the effect of high receptor densities and a saturable endocytic system. J Cell Biol 107: 801–810

Wu P-K & Park J-I (2015) MEK1/2 Inhibitors: Molecular Activity and Resistance Mechanisms. Semin Oncol 42: 849–862

Yao Z, Gao Y, Su W, Yaeger R, Tao J, Na N, Zhang Y, Zhang C, Rymar A, Tao A, et al (2019) RAF inhibitor PLX8394 selectively disrupts BRAF dimers and RAS-independent BRAF-mutant-driven signaling. Nat Med 25: 284–291

Yao Z, Torres NM, Tao A, Gao Y, Luo L, Li Q, de Stanchina E, Abdel-Wahab O, Solit DB, Poulikakos PI, et al (2015) BRAF Mutants Evade ERK-Dependent Feedback by Different Mechanisms that Determine Their Sensitivity to Pharmacologic Inhibition. Cancer Cell 28: 370–383

Yen I, Shanahan F, Lee J, Hong YS, Shin SJ, Moore AR, Sudhamsu J, Chang MT, Bae I, Dela Cruz D, et al (2021) ARAF mutations confer resistance to the RAF inhibitor belvarafenib in melanoma. Nature 594: 418–423

Yen I, Shanahan F, Merchant M, Orr C, Hunsaker T, Durk M, La H, Zhang X, Martin SE, Lin E, et al (2018) Pharmacological Induction of RAS-GTP Confers RAF Inhibitor Sensitivity in KRAS Mutant Tumors. Cancer Cell 34: 611–625.e7

Yin N, Ma W, Pei J, Ouyang Q, Tang C & Lai L (2014) Synergistic and Antagonistic Drug Combinations Depend on Network Topology. PLOS ONE 9: e93960

Yuan B, Shen C, Luna A, Korkut A, Marks DS, Ingraham J & Sander C (2020) CellBox: Interpretable Machine Learning for Perturbation Biology with Application to the Design of Cancer Combination Therapy. Cell Syst

Zhang C, Spevak W, Zhang Y, Burton EA, Ma Y, Habets G, Zhang J, Lin J, Ewing T, Matusow B, et al (2015) RAF inhibitors that evade paradoxical MAPK pathway activation. Nature 526: 583–586

